# RNA-specific local translation is patterned by condensates for multinucleate cell growth

**DOI:** 10.1101/2024.12.12.628219

**Authors:** Zachary M. Geisterfer, Ameya P. Jalihal, Sierra J. Cole, Amy S. Gladfelter

**Author notes:** These authors contributed equally (Zachary M. Geisterfer, Ameya P. Jalihal).

## Abstract

Coordination between growth and nuclear division is a common cell feature. In some syncytia, nuclei divide asynchronously throughout the cell but growth occurs only at discrete locations, raising the question how the processes are locally regulated and globally coordinated. In the syncytial fungus *Ashbya gossypii*, both cell-cycle progression and hyphal elongation require condensates formed by the protein Whi3 in complex with distinct mRNA species. We show that Whi3 condensates are enriched for translation regulators and are associated with local, spatially patterned translation of specific target RNAs near nuclei and growth sites. Whi3-RNA condensates can both promote and repress mRNA translation in an RNA- and condensate size-dependent manner in vitro. Condensate-interfaces are sites of translation, tunable by condensate composition, RNA valency and protein charge-state in vitro. Together, these data suggest that Whi3 condensates can generate a continuum of translation states that vary depending on the subcellular location and resident RNA sequences.

## Introduction

Large cells such as unicellular slime molds, filamentous fungi, alga, skeletal muscles and neurons operate on spatial scales at which diffusion becomes limiting for rapid, coordinated regulation at distal sites^1^. These cells face a challenge of mounting local responses such as growth and chemotaxis in coordination with global physiological requirements or developmental programs. This local-global challenge is exemplified in the growth of the multinucleate filamentous fungus *Ashbya gossypii,* in which the rate of nuclear division is coupled to the rate of hyphal growth to maintain a constant nucleus number-to-cytoplasm ratio^2^. However, nuclear divisions are often 100s of micrometers away from a growth site, raising the question as to how these processes are locally controlled and yet coordinated.

Previous work from our lab has shown that a condensate-forming RNA-binding protein called Whi3 is critical for patterning both nuclear divisions and polarized growth, implicating it as a potential point of coordination between these distinct processes^3,4^. Biomolecular condensates can both locally control biochemistry and be responsive to global signaling, serving as an attractive mechanism for coupling global cell states to a local process^5–7^. Condensates are controlled by post-translational modifications or direct environmental sensitivities that result in altered molecular interactions^8–12^. The strength, valency and specificity of interactions amongst components within the condensed phase can in turn impact its composition and physiochemical properties such as pore size, charge and pH^13–16^. This enables a condensate of specific molecular composition to dynamically change its properties depending on subcellular location or cell status.

A long-appreciated function of RNA-protein (RNP) clusters and condensates is to position and regulate mRNA translation. For example, stress granules and P-bodies are enriched for non-translating RNAs^17,18^. During embryogenesis, P granules concentrate and regulate mRNAs for germline specification^19,20^ and FXR1 granules show selective translation activation during spermiogenesis^21^. A series of recent studies uncovered the ability of Drosophila germline granules to locally promote translation of specific RNAs, indicating that these condensates can enhance translation^22–24^. RNP granules are also key passengers for long-range transport of repressed mRNAs in neurons for ultimate translation at distal processes in response to various cues^6,25,26^. Notably, across these varied contexts, RNP condensates must be able to switch between states where mRNAs are repressed and translated rather than the condensed state having a single function ^6,10,27,28^.

Our previous work showed that Whi3 condensates localize the G1 cyclin mRNA *CLN3* which is clustered near nuclei, and the formin mRNA *BNI1* which is enriched at hyphal tips^3,4,29^. Notably, these mRNAs encode proteins whose abundance and localization are tightly controlled: Cln3 shows extremely low abundance and is remarkably hard to detect across species^30^ and Bni1 shows a highly constrained localization to hyphal tips^31^, consistent with tight post-transcriptional control of their gene expression. Whi3 contains a glutamine rich region (QRR) that is critical for condensation, and an RRM-type RNA binding domain that binds a conserved RNA motif^3^. Whi3 is phosphorylated at various sites in response to stresses as well as growth-associated and cell-cycle signals^5^. Perturbing Whi3’s QRR or Whi3’s ability to bind mRNA both decrease Whi3 condensates and mRNA clustering in the cell^3^. Mutants with loss of mRNA spatial patterning have dramatically altered growth, nuclear division asynchrony and branching^3,4^. Here, we set out to understand the molecular functions of Whi3 condensates that contribute to these phenotypes.

We found that Whi3 condensate size and number correlate with hyphal growth rate and nuclear cycle progression. Motivated by the observation that the Whi3 interactome is enriched for regulators of translation and RNA metabolism, we used three orthogonal approaches to test the hypothesis that Whi3 condensates regulate the translation of target mRNAs in space. We first used a FISH-based strategy to probe spatial variation in ribosome association with Whi3 target mRNAs in cells. This revealed that *CLN3* and *BNI1* both show distinct spatial patterns of translation that are associated with Whi3 condensates. To test if Whi3 protein and target mRNAs were sufficient to regulate translation, we turned to two in vitro assays and observed that RNA sequence, number of Whi3 binding sites, condensate size and Whi3 concentration can all alter the translation outcomes in a minimal system. Finally, we used an imaging-based approach involving a real-time translation readout and observed a striking enrichment of translation within the dense phase and at the condensate interface. Together, these lines of evidence point to the ability of Whi3-RNA complexes to both activate and repress *CLN3* and *BNI1* translation depending on cell location and condensate properties. We propose that translation state-tuning allows Whi3 condensates to regulate local protein abundance of these critical regulators of cell cycle and growth and speculate this is a mechanism to integrate these processes.

## Results

### Whi3 condensates are spatially correlated with hyphal growth and cell cycle states

We previously reported that Whi3 forms spatially distinct condensates that localize G1 cyclin mRNA *CLN3* near nuclei, and formin mRNA *BNI1* to hyphal tips^3,4^. To visualize how Whi3 condensate localization relates to cell growth we performed 3D time-lapse fluorescence imaging of Whi3:tdTomato using lattice light sheet microscopy. We observed a surprising negative correlation between the tip elongation rate and the presence and proximity of Whi3 condensates to the hyphal tip (**Figure 1A, B**). We also measured hyphal elongation rates in mutants that were previously reported to decrease condensation including a mutant lacking the Whi3 glutamine rich region (QRR), and a phosphomimic of the conserved PKA phospho-site S637^3,5,32^. We confirmed previous observations of a strong reduction in tip condensates in the two mutant strains (**Figure 1C, left**) and found that both strains showed significantly faster hyphal elongation rates with decreased tip condensates (**Figure 1C, right**) than wildtype. These data show that the presence of a Whi3 condensate in the hyphal tip is negatively associated with tip elongation rate.

**Figure 1.**
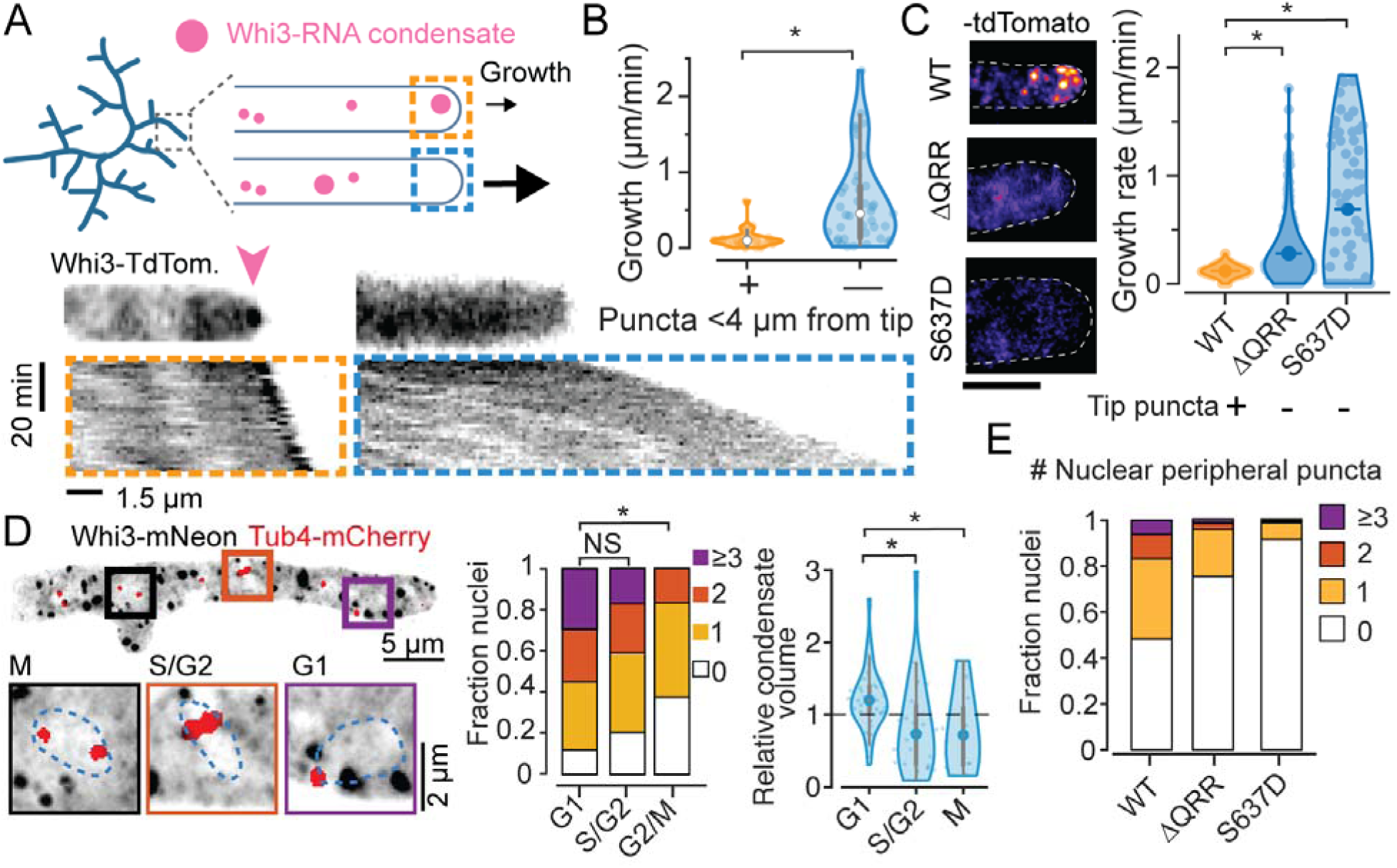
Whi3 condensate variability at hyphal tips and around nuclei is correlated with variability in hyphal elongation and cell cycle state. A. Schematic illustration of Whi3 condensates and growing hyphal tips in *Ashbya gossypii* (top) and representative timelapse kymographs of hypha expressing Whi3:tdTomato (bottom). Kymographs were created from MIPs of 3D lightsheet timelapses. Orange box shows an example of a hypha with a prominent visible condensate at tip, marked by a pink arow, and the blue box shows an example of a hypha without a large tip-proximal condensate. Scale bar x: 1.5 µm, time: 20 min, origin on top left. B. Instantaneous hyphal growth extracted from a total of 21 movies, split into 40 clips with or without tip proximal condensate (<4 µm from tip) each. * = p<0.01 using KS test. C. Representative images of WT Whi3:mNeon, non-condensate-forming *whi3-QRR*:tdTomato and *whi3 S637D:tdTomato Ashbya* hyphae (left) and hyphal elongation rates (right) measured using ConA pulse labeling. N>40 hyphae from >10 cells per strain. Scale bar 5 µm. D. Representative MIP of Tub4-mCherry expressing plasmidic Whi3-mNeon, scale bar 5 µm. Insets shows G1, S/G2 and G2/M nuclei with variable numbers of nuclear-peripheral Whi3 puncta, scale bar 2 µm. Stacked histograms quantifying relative nuclear proportions having varying peri-nuclear Whi3 puncta counts pooled from 3 biological replicates from a total of 238 G1, 88 S/G2, 24 G2/M phase nuclei. Violin plot of normalized condensate volume distribution binned by cell cycle state, of condensates lying within 1 µm of nucleus centroids. Volumes were normalized to the median volume within each hypha, from a total of 42 hyphae. * = p<0.01 using KS test. E. Stacked histograms quantifying relative nuclear proportions having varying peri-nuclear Whi3 puncta counts, from WT Whi3-tdTomato, n=180, *whi3*(ΔQRR):tdTomato, n=148, *whi3*(S637D):tdTomato, n=194.

We next used high-resolution confocal reconstructions of Whi3-mNeon/Tub4-mCherry cells to determine the distribution of Whi3 puncta around nuclei, where they were previously shown to associate with the ER membrane^29^. The periphery of nuclei, defined as the region 500 nm around to the edge of a segmented nuclear boundary, showed considerable variability in the number and size of Whi3 puncta (representative image, insets and histogram in **Figure 1D).** This size variability had clear cell cycle association based on spindle pole appearance which was visualized using Tub4-mCherry^33^. Nuclei with two well separated spindle poles, indicating late G2/M phases, had fewer and smaller peri-nuclear condensates relative to both G1 and S/G2 nuclei (**Figure 1D; right**). Mutant cells *whi3*(ΔQRR) and *whi3*(S637D) both showed decreased condensate number and size, leading to few nuclei with any proximal condensate and less variability amongst nuclei overall (**Figure 1E**). This is consistent with the increase in nuclear division synchrony phenotype observed in these mutants^4,5^. Thus, Whi3 condensate size, number and presence correlate with both hyphal growth and cell cycle state.

### Whi3 protein associates with regulators of mRNA and translation

To determine molecular functions for Whi3 condensates, we identified the proteins that associate with Whi3 using an affinity purification protocol to enrich and isolate Whi3-associated proteins. Whi3 condensates did not remain intact using standard IP-protocols, prompting us to design an alternative approach to enrich the condensed state of Whi3. Briefly, lipid-coated beads functionalized with Ni-NTA lipids were coated in recombinantly expressed 6xHis-Whi3. These beads recruit endogenously expressed Whi3-tdTomato from *Ashbya* extracts to the bead surface (**Figure 2A,B**). A negative control of beads coated only in Ni-NTA lipids was used to control for non-specific binding to lipids or and these beads did not recruit endogenous Whi3 from extracts (**Supplementary Tables 1-2**). Bead-enriched proteins were analyzed by mass spectrometry (**Figure 2A,C**). Among the significantly enriched proteins, we identified ribosome components, translation regulators and RNA decay factors (**Figure 2C; Supplementary Tables 1-2**). Notably, a majority of the proteins identified in these interactions are RNA-dependent (**Figure 2C**; 160 enriched proteins identified in non-treated lysates, 6 proteins identified RNAse A treated lysates). Thus, Whi3-RNA assemblies can associate with post-transcriptional regulators of RNA stability and protein translation, potentially acting as a scaffold for or a direct regulator of these processes.

**Figure 2.**
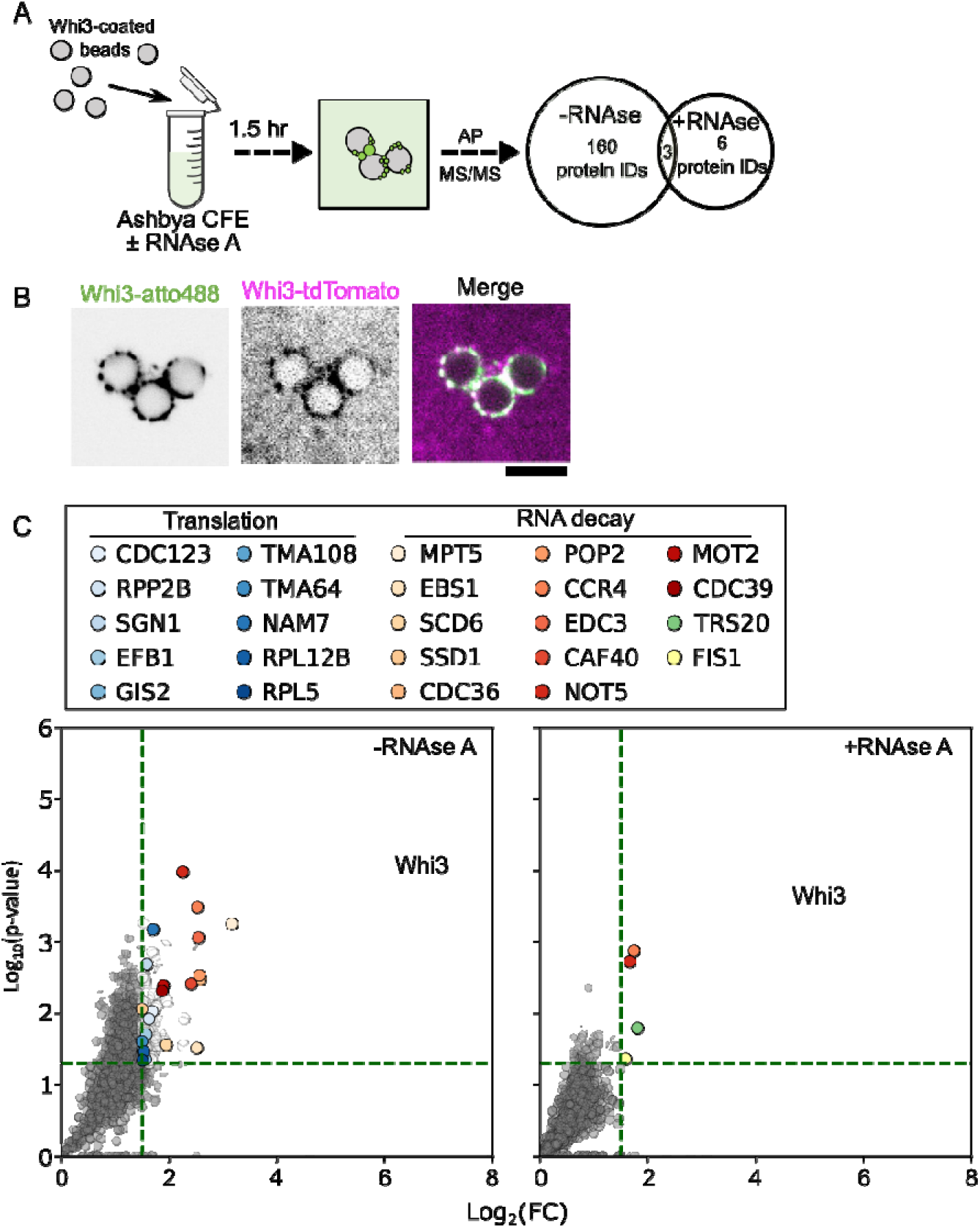
Whi3 condensates interact with regulators of translation and RNA metabolism. A. Whi3-assemblies can be purified from *Ashbya* cell-free extracts. Proteins identified are expressed as a Venn diagram indicating the number of enriched proteins for each treatment group. The central overlap indicates the number of unique protein IDs shared between the treatments. B. Fluorescence images showing microspheres coated with recombinantly expressed Whi3-atto488 recruit endogenously expressed Whi3-tdTomato from *Ashbya* lysates. Scale bar 5 µm. C. Protein IDs corresponding to mass spectrometry identification plotted as a function of fold-change (FC) over the control condition (microspheres without recombinant Whi3) for lysates with RNA intact (RNAse-) or treated with RNAse (RNAse+). Green lines indicate the criteria for significance. Translation regulators are indicated in light-blue and factors involved in RNA decay are shown in red. The closest yeast homolog is indicated above. Results are from three independently prepared *Ashbya* lysates (see methods).

### Whi3 target mRNAs show spatially variable translation

To examine if Whi3 condensates regulate translation in vivo we first performed polysome profiling on WT and *whi3*(ΔQRR). We reasoned that if Whi3 condensates influenced translation of its targets, the absence of condensates would result in a difference in polysome association of Whi3 targets *CLN3* and *BNI1*. Both RNAs showed polysome association (fractions 7-11) however only *CLN3* showed a detectable increase in a translating fraction and loss of a non-translating population in the *whi3*(ΔQRR) relative to WT (fractions 4-5) (**Figure S1A**). This result suggests that Whi3 condensates are associated with partial translation repression for *CLN3*, but this was not detectable for *BNI1*. However, polysome profiling is a bulk measurement and therefore insensitive to small local variations which are likely relevant for *BNI1* where hyphal tips represent a small fraction of total cell area.

We therefore next tested the hypothesis that Whi3 targets show spatial variation in translation in cells. We turned to the recently reported “tri-probe” proximity ligation (PL)-based strategy to detect ribosome associations with endogenous mRNAs^34^. Briefly, four and five sets of “padlock” and “primer” DNA probes tiling *CLN3* and *BNI1* respectively were used in conjunction with five Ag18s rRNA splint probes. The splint probes allowed ligation of proximal padlock probes by T4 ligase, creating a circular ssDNA template for rolling circle amplification (RCA) by Phi29 DNA polymerase. Rolling circle amplicons were detected via AlexaFluor647- or AlexaFluor555-labeled readout probes (**Figure 3A**). The method was adapted to *Ashbya*, and we observed bright, punctate PL-RCA signal that robustly colocalized with smFISH spots. On average, >50% of *CLN3* smFISH spots were colocalized with PL-RCA signals, whereas >80% of PL-RCA puncta overlapped with an smFISH spot, suggesting high specificity of detection (**Figure 3B**). We validated the sensitivity of this approach by perturbing translation rates pharmacologically with translation elongation inhibitors G418 or initiation inhibitor rapamycin^35,36^ which significantly decreased the number and brightness of PL-RCA puncta (**Figure 3C**). We conclude that PL-RCA puncta (henceforth “ribo”) represent not just ribosome proximity but active translation in *Ashbya*.

**Figure 3.**
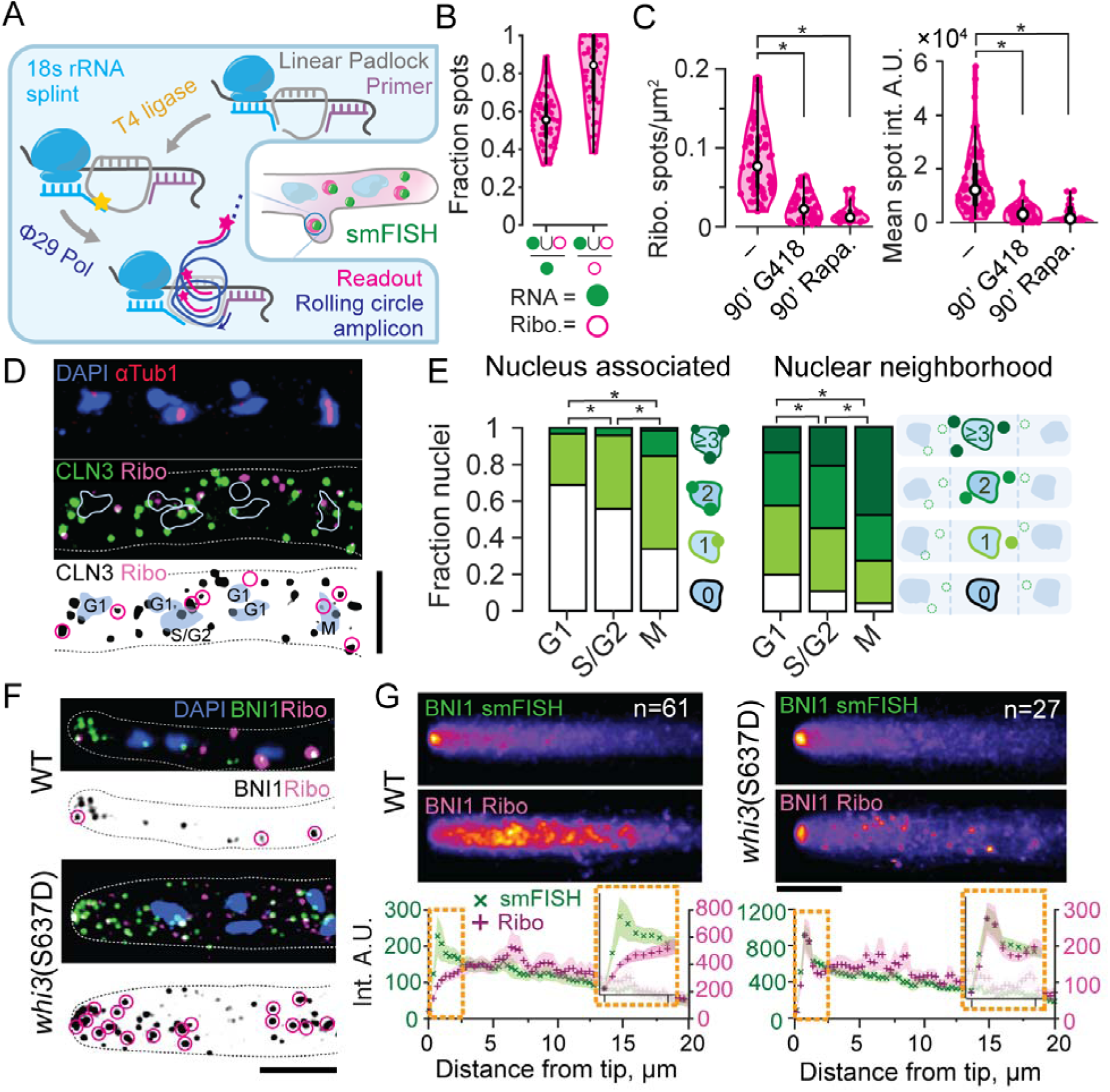
Whi3 target mRNAs show spatially biased translation in *Ashbya*. A. Schematic of tri-probe PL-RCA (“ribo”) strategy for detection of ribosome association using FISH in *Ashbya*. B. Most PL-RCA amplicons for *CLN3* show specific colocalization with *CLN3* smFISH spots, representing a majority of RNA spots. Spot masks in the smFISH (green filled circle) and RCA readout (magenta hollow circle) channels obtained using a semi-automated Otsu threshold were used to determine the fraction of colocalized spots with respect to counts of smFISH and PL-RCA spots per hypha, n=62 cells. C. PL-RCA amplicons represent true translation. The PL-RCA spot density per hyphal mask and mean brightness for *CLN3* both significantly decreased (KS test, p<0.01), upon 90 minutes of 200 µg/mL G418 (44 hyphae, 3757 RNA, 957 PL-RCA spots) or 90 minutes of 10 µg/mL rapamycin (17 hyphae, 348 RNA, 791 PL-RCA spots). Drugs were added to 10 mL of 14 h old cultures just prior to fixation. D. Representative image of CLN3 PL-RCA showing nuclear proximal translation. Blue, Hoechst-stained nuclei; red, AlexaFluor488 labeled secondary, for anti-Tub1 antibody; green, *CLN3* smFISH using TAMRA-labeled probes from Stellaris; magenta, CLN3-PL-RCA amplicons detected using AlexaFluor647-labeled readout probe, hyphae are outlined with dotted line. Bottom panel: schematic with nuclei in blue and their cell cycle state indicated, black represents *CLN3,* magenta circles indicate translating RNAs. Scale bar 1 µm. E. Cell cycle stage correlations with counts of *CLN3* Ribo at the nuclear-periphery (left) or nuclear-neighborhood (right). *=p<0.01, KS test on average spot count per nucleus. F. Representative images of hyphal tips with *BNI1* smFISH (green) and Ribo (magenta) in WT and *whi3*(S637D). Scale bar 5 µm. Lower panels show identified BNI1 RNAs in black, translation in magenta circles. G. Mean projection images of MIPs of *BNI1* smFISH (top) and Ribo (bottom) channels across hyphal tips for WT (61 tips) and *whi3*(S637D) (27 tips) . Mean intensity and SEM at points sampled at 0.365 µm interval along the length of the hyphae aligned to the tip. Orange boxes show magnified insets of the profiles 2.5 µm closest to the hyphal tip. Scale bar 5 µm.

We first applied this method to study spatial variation in translation of *CLN3* mRNA, which was previously reported to form clusters near nuclei in a Whi3-dependent manner^3^. We observed a clear pattern of nuclear-periphery-associated *CLN3*-ribo spots (**Figure 3D**). Based on data in **Figure 1D**, we next combined *CLN3* PL-RCA with immunofluorescence for tubulin to determine local cell cycle state of translation sites. M-phase nuclei showed the greatest number of nuclear-peripheral as well as nuclear proximal *CLN3* translation spots, followed by S/G2, then G1, which showed the fewest spots (**Figure 3E**). Importantly, these data show that the Whi3 target mRNA *CLN3* is locally translated in specific windows of the cell cycle.

We next investigated the sites of translation of *BNI1*, whose mRNA and protein were previously reported to cluster at hyphal tips in a Whi3-binding-dependent manner^4,31^. While smFISH revealed the expected clustering of *BNI1* mRNAs at tips, we saw infrequent sites of translation at these locations. We considered that the lack of signal could be an artifact of the growing hyphal tip being crowded with secretory vesicles^31^, making it inaccessible to either translation machinery and/or the RCA probes. However, *whi3*(S637D), which has rapid tip growth and small Whi3 puncta (**Figure 1C**; right), showed both *BNI1* mRNA clusters and enriched translation of those mRNAs at the hyphal tip (**Figure 3F-G**). While RNA abundance at the tip differs between these strains, the change in *BNI1* translation pattern suggests Whi3 influences the spatial localization of *BNI1* translation in WT. We conclude that *BNI1* has variable translation at hyphal tips but is predominantly repressed at tips in WT cells. Thus, both Whi3 target mRNAs examined show spatially and temporally variable translation.

### Whi3 condensates can be associated with sites of translation in vivo

To determine the relationship between the translation of Whi3 target mRNAs and condensates in vivo, we repeated PL-RCA and smFISH simultaneously with immunofluorescence against Whi3. As these are fixed cells, we are defining Whi3 condensates as puncta that are brighter than background signal and localized in the same areas where Whi3 puncta fusion and turnover have been previously analyzed^37^. We observed many but not all translation events associated with Whi3 puncta (**Figure S2A,B** insets). The spatial distribution of condensate-associated translation spots was measured by calculating Whi3 signal at translation spots and comparing this to Whi3 signal associated with non-translating RNAs. This revealed that *CLN3* translation was specifically enriched at Whi3 puncta at the nuclear periphery (0.66 µm from the nucleus centroid) when compared to the rest of the cytoplasm (**Figure 4A,C**, more examples in **Figure S2C**). Similarly, this effect does not arise from random chance based on the number of pixels occupied by Whi3 signal **(Figure 4C**; top panel). *BNI1* was overall infrequently translated at tips, but when *BNI1* translation was detected at tips, it is often associated with a Whi3 puncta when compared to translating *BNI1* mRNAs away from the tip in the same hypha (**Figure 4B,D**). Thus, we see that subsets of Whi3 condensates near nuclei (for *CLN3*) and at tips (for *BNI1),* are associated with local translation of these RNAs.

**Figure 4.**
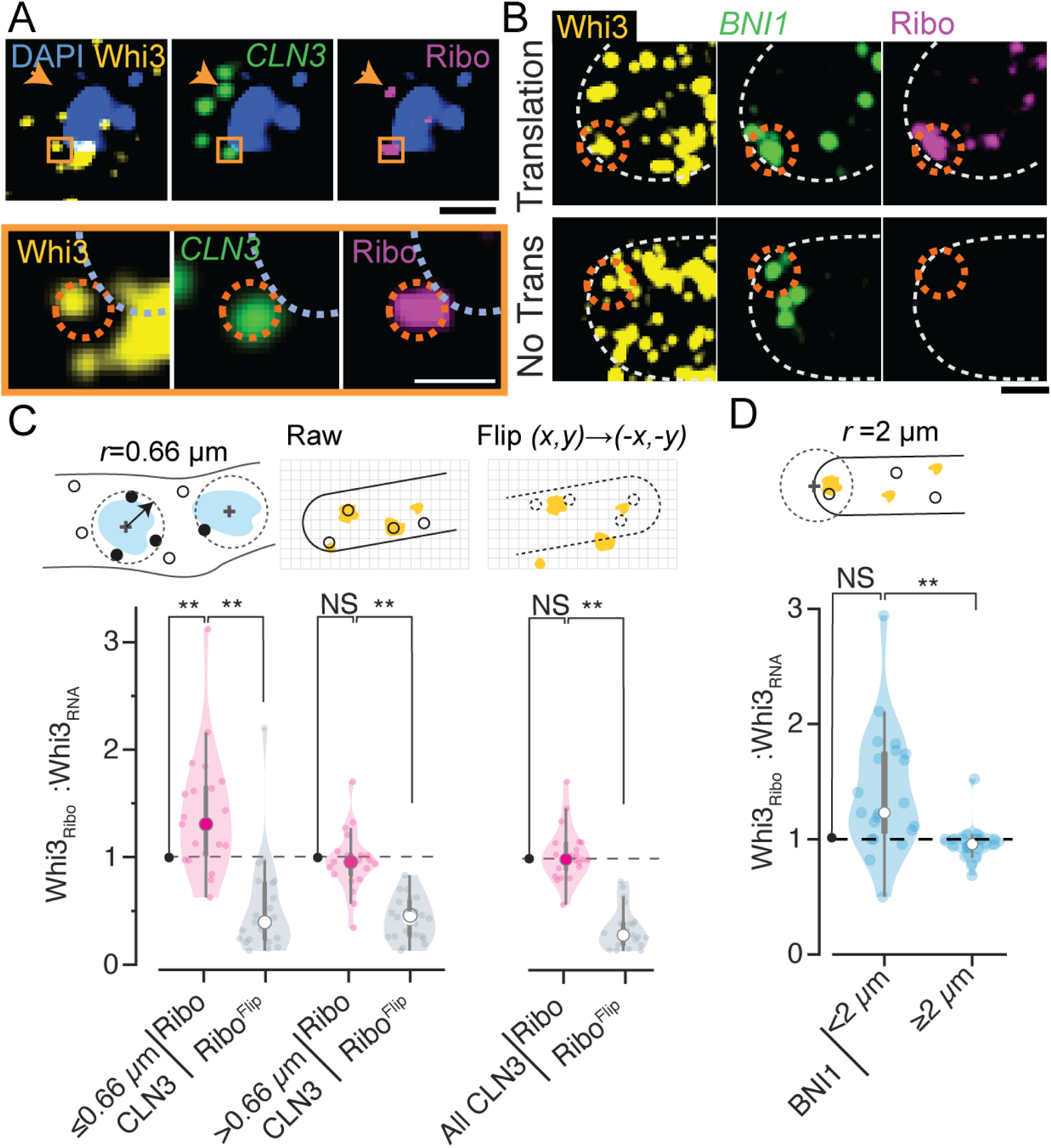
Condensate associated translation is spatially patterned in vivo. Whi3-mNeon IF puncta colocalize with translating *CLN3* and *BNI1* at nuclei and tips. Representative 4-color fluorescence images of Whi3 condensate in conjunction with smFISH and PL-RCA for Whi3 targets. A. Representative image of *CLN3* translation at nuclear periphery. Orange arrow marks a translating *CLN3* RNA spot away from the nucleus and not associated with a condensate, and the orange box, shown in insets, highlights a translating RNA spot on the nuclear periphery associated with a condensate. Scale bar: image, 1 µm, the inset, 0.5 µm. Also see Figure S2C. B. Representative image of *BNI1* associated with condensates at hyphal tips. Condensate-associated RNAs can translate (top) or be repressed (bottom). C. Top panel: schematic showing nuclear proximity analysis and spatial randomization by flipping Ribo channel (black circles) and measuring masked Whi3 channel (yellow). Bottom row: Median Whi3-mNeon fluorescence intensity associated with Ribo-associated RNA spots relative to median Whi3-mNeon intensity associated with all *CLN3* RNA spots per hypha. Mean Whi3 intensities are compared to a spatially randomized “flipped” image. *CLN3* translation is enriched at higher Whi3 intensities near the nuclear periphery (0.66 µm from centroid) relative to cytoplasmic translation, which is indistinguishable from median RNA-associated Whi3. D. Relative Whi3-mNeon fluorescence intensity associated with *BNI1* Ribo spots 2 µm from hyphal tips relative to spots away from tip per hypha. Relative intensities calculated across N=21 hyphae for *CLN3*, and N=38 hyphae for *BNI1*. ** indicates p<0.01 for t-test on population means or mean relative to value 1.

### Intrinsic features of Whi3 condensate components can modulate translation

To determine if Whi3-dependent translation was intrinsic to the condensates or required other cell inputs, we turned to in vitro translation assays. First, we performed a bulk in vitro luciferase reporter assay in which Nanoluciferase (*Nluc)* CDS was placed downstream of 5’UTRs consisting of *CLN3* or *BNI1* sequences containing Whi3 binding sites (**Figure S3A**).

Recombinant Whi3 protein and in vitro transcribed reporter RNAs were mixed prior to the addition of Rabbit Reticulocyte (RR) lysate, and end point nano-luciferase activity was measured as a function of Whi3 concentrations. Two Whi3 concentrations corresponding to the dilute phase of the *Ashbya* cytoplasm (∼50 nM)^38^, and two concentrations that resulted in the formation of Whi3 condensates *in vitro* were tested (**Figure 5A-D**). We observe little change in luciferase activity compared to the no Whi3 control for either *CLN3-Nluc* or *BNI1-Nluc* in the 1-phase regime (**Figure 5A,C**) suggesting the Whi3 does not contribute substantially to promote translation in small complexes expected in these concentrations below the C_sat_ for phase separation. At condensate-forming Whi3 concentrations (1 and 3 µM), translation of *CLN3-Nluc* showed monotonic Whi3 dose-dependent repression of up to >100-fold repression at 3 µM (**Figure 5A,B**). The substantial levels of repression observed for *CLN3-Nluc* were not driven by RNA turnover, as RNA degradation was the same +/- Whi3 (**Figure S3B**). In contrast, condensate-promoting Whi3 concentrations showed a biphasic response in *BNI1-Nluc* translation: 1 µM Whi3 protein showed >2-fold enhancement of translation while 3 µM showed strong repression (**Figure 5C,D**). These data show the intrinsic properties of Whi3 and its target RNAs are sufficient to produce some concentration-dependent heterogeneity of translational outputs in a simplified in vitro system.

**Figure 5.**
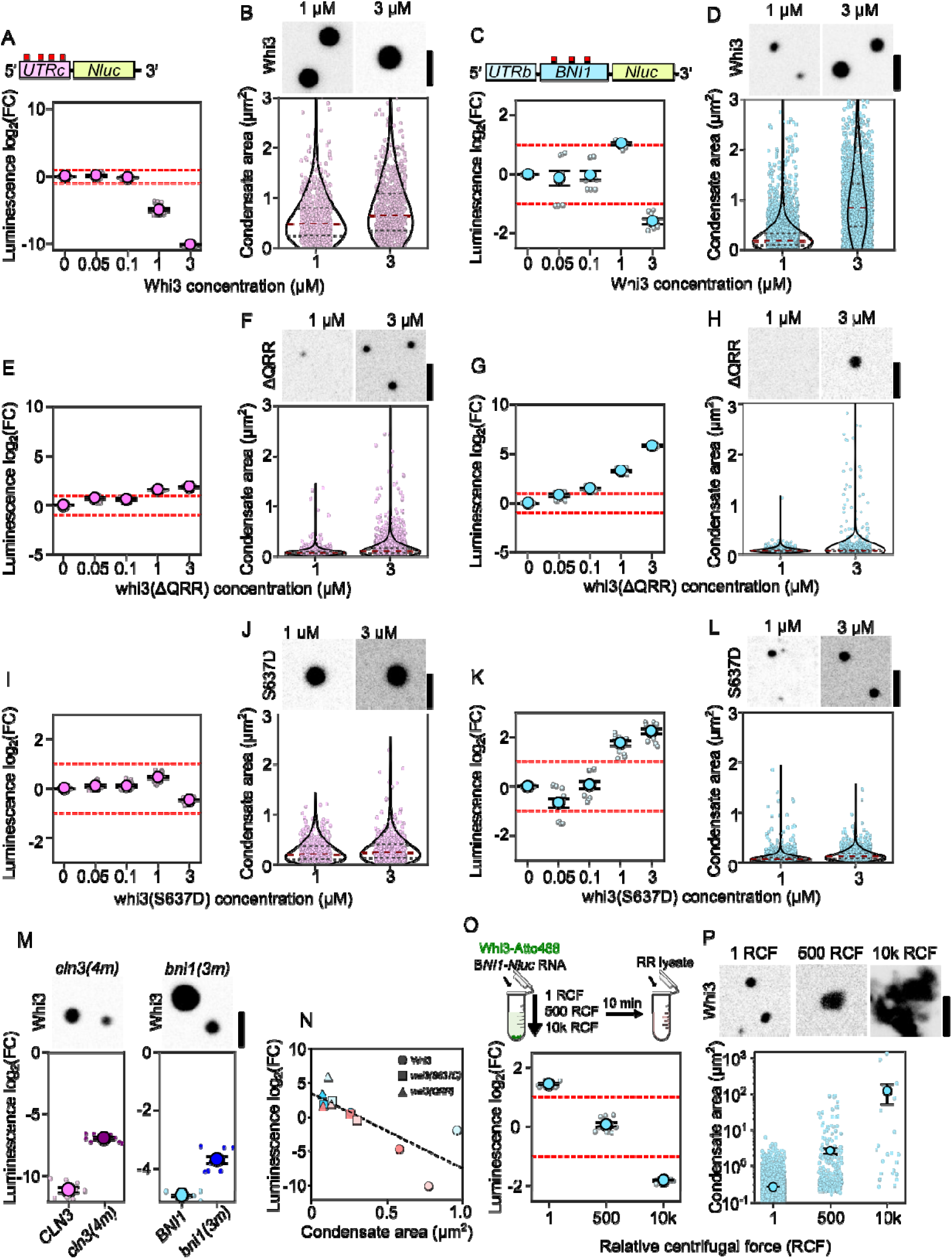
Condensate-promoting interactions promote translation repression in vitro. Luciferase reporters of Whi3-target RNAs show distinct Whi3-dependent translation. In **A,C,E,G,I,K** luciferase activity is displayed as log_2_(fold change) over no-Whi3 conditions for a Whi3 dose response. Dashed red lines indicate log_2_(fold change) = 1. Measurements are taken from three separate RR lysates with three technical replicates per lysate (see Methods). Means are displayed +/- SEM. Representative images of the respective condensates are shown in **B,D,F,H,J,L**, along with violin plots showing condensate size distributions. Each dot represents a condensate and measurements were taken from reactions prepared with three different RR lysates. Scale bar = 2.5 µm. The reporter assays contain: **A,B**. *CLN3-Nluc,* WT Whi3; 1 µM (n=792) or 3 µM (n=900) condensates. **C,D**. *BNI1-Nluc,* WT Whi3, 1 µM (n=5201) or 3 µM (n=1890) condensates. **E,F.** *CLN3-Nluc,* whi3(ΔQRR); 1 µM (n=1310) or 3 µM (n=3985) condensates. **G,H**. *BNI1-Nluc,* whi3(ΔQRR); 1 µM (n=785) or 3 µM (n=1718) condensates. **I.J**, *CLN3-Nluc,* whi3(S637D); 1 µM (n=593) or 3 µM (n=607) condensates. **K,L** *BNI1-Nluc,* whi3(S637D); 1 µM (n=3449) or 3 µM (n=2020) condensates. **M**. Representative images of Whi3 condensates formed with either *cln3(4m*) or *bni1(3m*). Mutant reporter RNAs *bni1(3m)-Nluc* and *cln3(4m)-Nluc* were designed to preserve nucleotide composition of Whi3 binding sites but scramble the Whi3 recognition motifs. Luminescence values are displayed as log_2_(FC) over no-Whi3 conditions. Measurements are taken from three separate RR lysates with three technical replicates per lysate (see Methods). Means are displayed +/- SEM. Scale bar = 2.5 µm. **N.** Luciferase activity from all condensate-forming reactions plotted as a function of condensate area. Whi3 displayed as circles, whi3(S637D) as squares and whi3(QRR) as triangles. *CLN3-Nluc* is displayed in pink and *BNI1-Nluc* in blue, with lighter and darker fill indicating 3 and 1 µM conditions respectively. **O**. Translation of *BNI1-Nluc* in reactions containing 1 µM Whi3 left to form condensates on the bench top (1 RCF) or centrifuged at 500 or 10000 RCF prior to the addition of RR lysate. Luminescence values are shown as log_2_(FC) over RNA-only conditions spun at 1, 500, or 10000 RCF respectively. Measurements are taken from three separate RR lysates with three technical replicates per lysate (see Methods). Means are displayed +/- SEM. **P**. Fluorescence images of Whi3 condensates left to form on the bench top (1 rcf) or under 500 or 10000 rcf. Scale bar = 2.5 µm. Condensate area of *BNI1-Nluc* condensates 1 µM Whi3 conditions left to incubate on the bench top (n=5201), at 500 rcf (n=389), or 10000 rcf µM (n=23). Individual dots indicate a single condensate. Measurements were taken from reactions prepared with three different RR lysates.

While different concentration regimes could recapitulate enhanced and repressed translation of *BNI1*, the enhancement of *CLN3* translation as seen in a subset of nuclei in cells was not captured in vitro. We hypothesized that modulation of translation by condensates in cells could arise from changes in the intermolecular interactions that drive and maintain condensates. To test this hypothesis, we measured translation in reactions supplemented with whi3(ΔQRR) to represent weakened protein multivalency and with whi3(S637D) to represent phospho-regulation ^3,4,39^. Whi3(ΔQRR) at low concentrations (50 and 100 nM) resulted in modest increases in RNA translation for both reporter RNAs suggesting some small assemblies or nanoclusters can enhance translation in the context of a weakened protein-protein interactions (**Figure 5E,G**). Higher concentrations of whi3(ΔQRR) generate smaller condensates compared to similar concentrations of WT Whi3 for both *CLN3*- and *BNI1-Nluc* as anticipated, (**Figure 5F,H** compared to **Figure 5B,D**) and resulted in pronounced enhancement of translation for both RNAs relative to the low concentrations regimes and the no-Whi3 control (**Figure 5E,G**).

Concentrations of whi3(S637D) in the 1-phase regime resulted in no changes in translation for either *CLN3-Nluc or BNI1-Nluc* (**Figure 5I,K**). At higher concentrations, whi3(S637D) forms smaller condensates with both *BNI1-* and *CLN3-Nluc* mRNAs relative to WT Whi3 (**Figure 5I-L**) and distinct changes to translation. We observed a loss of translation repression for *CLN3* while *BNI1* experienced an enhancement in translation (**Figure 5I,K**). These results are also consistent with our observations in vivo, where cells expressing whi3(S637D) tend to form fewer and smaller Whi3 condensates and experience enhanced *BNI1* translation at tips. Notably, this change in translation regulation appears to originate from condensate properties and not RNA binding affinity in the mutant (**Figure S6A**). Finally, we also modulated the valence of the system by altering protein-RNA interactions by mutating Whi3 binding sites on *CLN3* and *BNI1* ^37,38,40^. At 3 µM WT Whi3, both mutant RNAs still formed detectable condensates, presumably due to non-specific RNA-protein interactions, however both RNAs had decreased translation repression relative to RNAs with intact binding sites (**Figure 5M**).

Notably, across all the luciferase assays, the highest translational repression for both *CLN3-Nluc* and *BNI1-Nluc* were seen in conditions that give rise to larger condensates (**Fig 5N**). However, Whi3 concentrations that drive the formation of large condensates also result in a substantial dilute phase concentration (**Figure S3D,E**), which potentially may repress translation even in the absence of visible condensates (**Figure S3C**), making it difficult to definitively say which phase dominates the translation regulation. To address this, we forced condensates to merge via centrifugation without changing RNA and protein concentrations^40^ (**Figure 5O**; schematic). Importantly, RR lysate was added directly to the reaction after centrifugation. Increasing centrifugation force resulted in progressively larger condensates (**Figure 5P**) that corresponded with increasing translation repression (**Figure 5O**). Since this centrifugation is not expected to shift the dense or dilute phase concentrations, this suggests that the size of individual condensates is relevant for repression. Thus, the mRNA and Whi3 protein components have an intrinsic dynamic range for repressing or promoting translation depending on concentration, condensate size and intermolecular interaction strengths.

### Whi3 condensates are sites of translation in vitro

The luciferase assay is a bulk measurement that combines many scales of assemblies spanning RNP complexes, nanoclusters and mesoscale condensates ^41^. To directly visualize the location of translation relative to Whi3 condensates *in vitro* we used the “moon tag” system^42^. MoonTag peptides, arising from RNA reporters containing repeats of the gp41 peptide (“MoonTags”) in the CDS^42^, bind fluorescently labeled gp41 nanobody with nanomolar affinity as they emerge from ribosomes, resulting in fluorescence signal that reports translation elongation (**Figure 6A**). We adapted this system for RR lysates, and generated *CLN3-* and *BNI1-MoonTag* reporter mRNAs (**Figure S4A**). Condensates were pre-formed by mixing Whi3 and *MoonTag* reporter mRNAs before supplementing the reaction with RR lysate (**Figure 6B**). Condensates formed with both RNA reporters showed frequent association with punctate MoonTag nanobody signal (**Figure 6C,D**). To check if the MoonTag signal represents recruitment or retention of previously translated nanobody-bound peptides to the condensate, RNAs were allowed to translate in the absence of Whi3 for 1 h with 500 nM Nb-mCherry. These lysates were treated with cycloheximide for 1 h before the addition of Whi3-condensates and an additional 500 nM Nb-mCherry. We saw a near-complete loss of MoonTag signal associated with condensates, comparable to cycloheximide-arrest of translation (**Figure 6F**), suggesting that any MoonTag signal in the condensate represents active translation. Notably, this >50-fold drop in condensate associated translation under cycloheximide-arrest only resulted in a modest increase in condensate size, suggesting Whi3 condensates are not enriched in stalled translation complexes (**Figure S4C**). Thus, Whi3 condensates can act as sites of translation in vitro, as was observed within cells (**Figure 4**). We next repeated the experiment in conditions that had led to less repression in the bulk luciferase assay including RNA valency mutants *cln3(4m)-* and *bni(3m)-MoonTag* reporters (**Figure 5M,N**), and phosphomimic, whi3(S637D) mutants (which only formed condensates with *CLN3* Moontag reporter). In all contexts, these mutants were associated with brighter MoonTag signal, despite condensates containing less RNA overall in the case of the RNA binding mutants (scatterplots in **Figure 6C,D, E)**. These results support that the dense phase of Whi3-RNA condensates can directly regulate translation, as suggested by both bulk luciferase assays (**Figure 5**) and experiments in cells (**Figure 4**).

**Figure 6.**
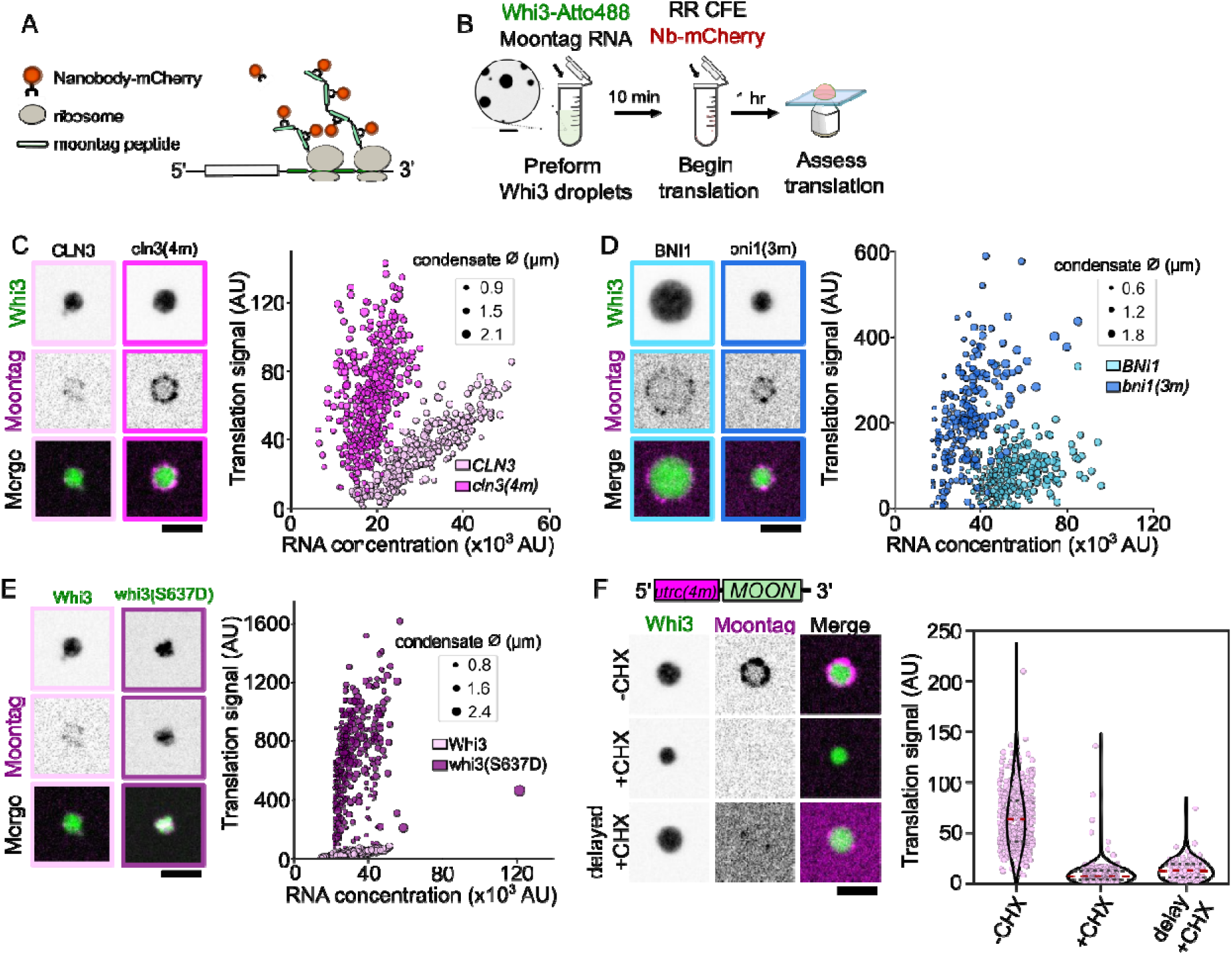
RNA valency and Whi3 phospho-state tune condensate-associated translation in vitro. **A.** Design of MoonTag reporter assay. **B.** Schematic of MoonTag reaction in cell lysates. **C.** Fluorescence image of a Whi3 condensates formed with either *CLN3*- or *cln3(4m)- Moon*, showing translation of mRNAs. Plots show translation signal as a function of RNA concentration for condensates formed with *CLN3-Moon* (light-pink; n=299) or *cln3(4m)- Moon* (dark-pink; n=460). Scatter points are calibrated by condensate diameter. **D.** Fluorescence image of a Whi3 condensate formed with either *BNI1*- or *bni1(3m)-Moon* showing translation of mRNAs. Plots show translation signal as a function of RNA concentration for condensates formed with either *BNI1-Moon* (light-blue; n=299) or *bni1(3m)-Moon* (dark blue; n=262). Scatter points are calibrated by condensate diameter. **E.** Fluorescence image of a whi3(S637D) condensates formed with *CLN3-Moon* showing translation of mRNAs. Plots show translation signal as a function of RNA concentration for condensates formed with *CLN3-Moon* and Whi3 (light-pink; n=299) or whi3(S637D) (purple; n=369). Scatter points are calibrated by condensate diameter. **F.** Fluorescence images showing translation of *cln3(4m)-Moon* at Whi3 condensates in RR lysates where translation is intact (-CHX), supplemented with cycloheximide (+CHX), or in lysates that have been allowed to translate *CLN3-Moon* mRNAs prior to either the addition of cycloheximide or Whi3 droplets (delayed +CHX). Scale bar = 2.5 µm. Quantification of average translation signal per condensate in translation competent lysates (-CHX; n=460), lysates supplemented with cycloheximide (+CHX; n=253), or lysates that have first been allowed to translate *CLN3(0xWBS)-Moon* prior to the addition of either cycloheximide or Whi3 droplets (delayed +CHX; n=127). Measurements were taken from reactions prepared with three different RR lysates.

### Translation is enriched at the interface of Whi3 condensates

Interestingly, the MoonTag signal observed at Whi3 condensates was not homogeneous across the condensate volume. While MoonTag signal was enriched within the condensate relative to the dilute, the signal was brightest at the condensate periphery (**Figure 7A,B**). Similarly, condensates formed with RNA valency mutant *cln3(4m)-* and *bni(3m)-MoonTag* also displayed annular localization of the MoonTag signal but it was significantly brighter (**Figure 6C,D**), and the peak of the signal showed a shift towards the interior of the condensate (**Figure 7C,D**; see MoonTag_Rmax_) compared to the wild-type sequences. The interface-enrichment of translation we observed does not reflect the position of RNAs within the condensates, as RNAs themselves showed a uniform localization profile across these condensates (**Figure S4B**). Finally, we probed the accessibility of the condensate interior with fluorescently labeled ribosomes.

**Figure 7.**
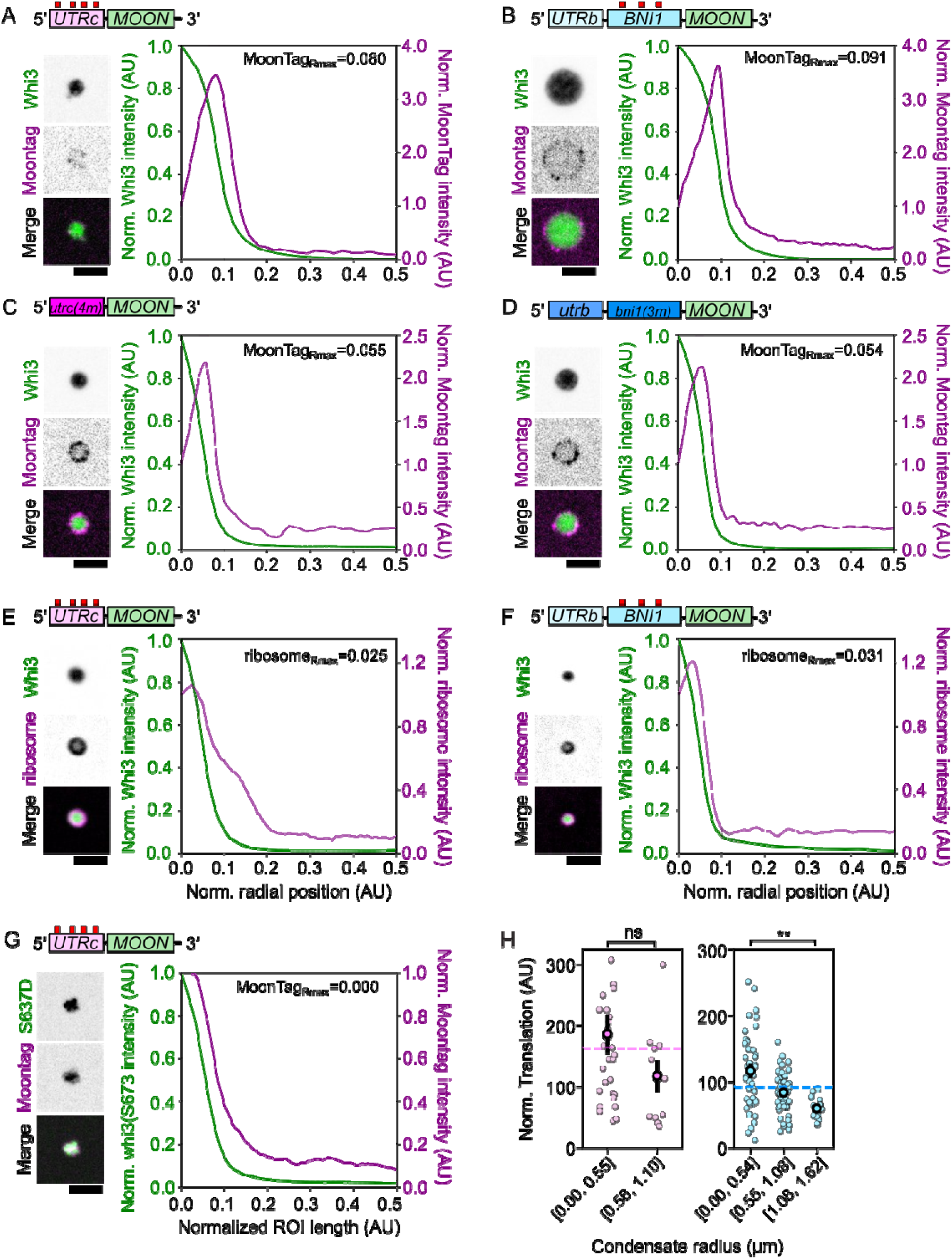
Condensate interfaces are sites for translation in vitro. Moontag reporter RNAs show a distinct interface-associated translation. In **A,B,C,D,E,F,G,** radial intensity profiles are displayed for Whi3 condensates formed with MoonTag reporter RNAs, showing fluorescence intensity as a function of radial position in the condensate for Whi3, MoonTag Nb-mCherry, or ribosomes. Dotted grey line marks normalized radius of the condensate, equal to 0.1 AU. The radial position associated with the maximum MoonTag or ribosome intensity recorded, MoonTag_Rmax_ or ribosome_Rmax_, is shown. Trace shows the average intensity ± SEM. Radial intensity measurements were acquired from condensates taken from three separate RR lysates. Representative images of the respective condensates are shown in **A,B,C,D,E,F,G.** Scale bar = 2.5 µm. Radial intensity profiles are displayed for condensates formed with: **A**. *CLN3-Nluc* and WT Whi3 (n=42). **B.** *BNI1-Nluc* and WT Whi3 (n=123). **C.** *cln3(4m)-Nluc* and WT Whi3 (n=147). **D**. *bni1(3m)-Nluc* and WT Whi3 (n=127). **E**. *CLN3-Nluc* and WT Whi3 (n=75). **F.** *BNI1-Nluc* and WT Whi3 (n=36). **G**. *CLN3-Nluc* and Whi3(S637D) (n=163). **H.** Average MoonTag signal of the condensate interface normalized by RNA concentration and plotted as a function of condensate radius. Each dot represents an individual condensate formed with either *CLN3-* (n=42) or *BNI1-Moon* (n=123), taken from at least three separate RR lysates. ** indicates P-value < 0.001 as measured by two-tailed t-test.

Interestingly, the condensate localization of ribosomes resembled the spatial bias we observed with translation, with the highest ribosome signal occurring at the perimeter (**Figure 7E,F**). The enrichment of both translation and ribosomes at the interface of condensates led us to wonder whether condensate-size-dependent effects on translation measured in bulk assays (**Figure 5N-P**) arise from smaller condensates having a larger surface area to volume. We measured the average intensity of the translation annulus associated with condensates (**Figure S5A**) and normalized this by the RNA concentrations at that position in the condensate. These measurements revealed that, in general, smaller condensates are associated with more translation than larger condensates (**Figure 7H**). These differences in translation-association could arise from smaller condensates having lower Whi3 concentrations (**Figure S5B**).

We next investigated if interface enrichment is a general feature of condensate-associated translation, given the recent work in Drosophila germ granules^22–24^. Condensates formed with whi3(S637D) had internal enrichment for MoonTag signal (**Figure 7G**), suggesting that changes in charge state of the protein can alter localization of translation to the interior. We next mixed our *CLN3-Moon* reporter RNA with two other well described phase separating RNA-binding proteins (RBPs), Fus and N-protein (**Figure S4D,E**). Fus condensates recruited MoonTag nanobody independent of translation, making them unsuitable to investigate translation (**Figure S4D**), while N-protein condensates completely repressed the translation of the reporter RNA (**Figure S4E**). The interface-enriched translation we observe for Whi3 condensates is therefore specific to particular forms of Whi3-target RNA condensates and controlled in part by the specific RNA-protein and protein-protein interactions therein.

Finally, to reconcile the condensate-associated translation seen with the Moontag system with the bulk repression seen in the luciferase assays of *CLN3* and the biphasic translation of *BNI1* in the two-phase regime, we turned to a minimal mass-balance model of translation arising from various RNA species in solution^43^. Overall translation is represented as a sum of contributions from unbound RNA, RNA bound to Whi3 in the dilute phase and RNA in the dense phase, and translation rates were estimated from luciferase data (**Supplemental methods**). These phase-specific translation rates, along with the inverse correlation between condensate size and translation were sufficient to recapitulate the qualitative features of bulk translation. The mass-balance model supports that differences in dense phase translation efficiency are sufficient to explain RNA-specific behavior at the phase boundary (**Fig 5A,D**). Condensate-specific translation can thus significantly shape bulk translation despite only constituting a small volume fraction of the solution in these systems.

## Discussion

In this study, we set out to examine the molecular function of Whi3 condensates in regulating nuclear division cycle and polarized cell growth. We found that condensates change size, number and presence in association with nuclear divisions and cell growth. These dynamics in condensate appearance were also associated with distinct subcellular translation states. *CLN3* mRNA is preferentially translated in association with nuclear-proximal Whi3 condensates in mitosis, when the condensates are smallest. *BNI1* translation is associated with condensates when it is detected at tips in wild-type cells. Both transcripts also show repression associated with Whi3, with large windows of the cell cycle and periods of tip growth without any protein production and specific conditions in vitro associated with repression. The balance of repression and activation can be modulated by mutations in either the protein or mRNAs that alter the intermolecular interactions and change condensate features. Both cell and in vitro data presented indicate that Whi3 condensates can both host and repress translation of resident mRNAs depending on context, molecular interactions and condensate composition.

This duality in function contrasts with contexts where condensates act as only an on or off switch where the condensed state is unifunctional. This gradation in function suggests a potential mechanism for metered production of target proteins, aligned with the cell cycle or growth requirements of the cell (**Figure 8**). This is akin to a switch between long-term storage followed by a punctuated translation as seen in germ granules, but Whi3 condensates appear to offer a pulsed local supply of proteins as required for oscillating nuclear division and polarized hyphal growth ^22–24^. This metered translation is associated with the ability to generate multiple co-existing polarity sites as mutants with continuous Bni1 translation lack lateral branches and instead grow from just a few dominant hyphal tips. Our in vitro assays show that the combination of translation repression and activation can manifest from a minimal system of the Whi3 protein and its target RNAs. While in our study we use mutants of Whi3/mRNAs, presumably inputs from cell signaling that similarly modulate protein-protein or protein-RNA interactions can shift between promoting and inhibitory states of Whi3. Whi3 condensates thus show a continuum of states to locally control translation.

**Figure 8.**
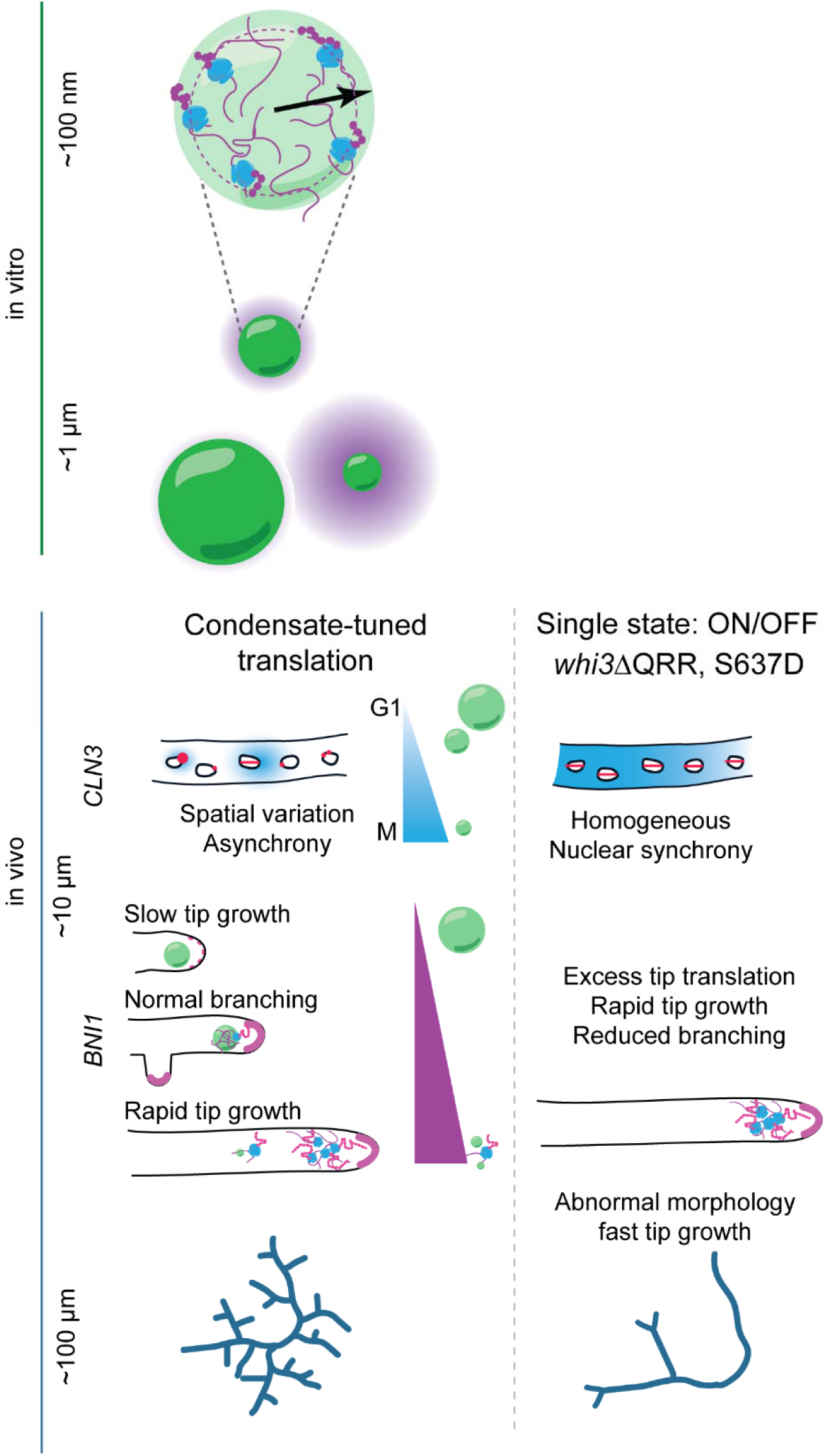
Whi3 condensates coordinate hyphal growth by dynamically switching translation state to tune mRNA-specific local translation. Top to bottom: 100 nm: RNA identity, binding-site number and Whi3 phospho-state shape protein translation at condensate interfaces. 1 µm: This results in a continuum of translation states that is additionally modulated by condensate size. 10 µm: This continuum is also reflected in the spatial variation in the translation of Whi3 targets at the cell scale in vivo. Condensates dynamically vary with cell cycle and growth rate resulting in a a range of translation states for *CLN3* and *BNI1* respectively. This is associated with nuclear division asynchrony and normal branching, along with variable tip elongation rates. However, the loss of condensate-promoting interactions also disrupts translation tuning, and results in nuclear division synchrony and poor branching, as a consequence of spatially homogeneous *CLN3* translation and excessive *BNI1* translation at tips. At the 100 µm scale, these differences results in dramatically different branching morphologies and growth rates.

Syncytial cells like filamentous fungi must coordinate large scale-physiological states with local responses such as polarized growth and nuclear division. In *Ashbya*, a shared condensate protein in complex with different sets of mRNAs acts in distinct local roles. Whi3 condensates both localize *CLN3* and tune translation at the nuclear periphery. *CLN3* shows an unexpected bias for translation at the periphery and neighborhoods of non-G1 phase nuclei, and substantial repression around G1 nuclei. The anti-correlation of translation with G1 phase was unexpected, given that *CLN3* encodes a G1 cyclin, but may reflect a feature of cell cycle regulation due to the syncytial nature of these cells. These differences in spatial translation may strongly shape the local abundance of this short-lived nuclear-localized protein that has unfortunately been challenging to directly detect in a variety of systems ^30,44^. We previously reported that Whi3 CDK phosphomutants show increased nuclear division synchrony ^5^. Given these same mutants show altered Whi3 condensates through the cell cycle, it is tempting to predict that CDK-regulation controls the size and abundance of Whi3 condensates in the vicinity of nuclei to pulse Cln3 production. It is intriguing that a G1 cyclin is translated by mitotic nuclei and we predict this may be a mechanism by which nuclei promote neighbors to be out of phase with each other.

Budding yeast, which also has a Whi3 homologue with a QRR (albeit substantially shorter than the one in *Ashbya*) has been shown to repress *CLN3* but there is no clear evidence of the condensate state activating translation and Cln3 translation is highest in G1^45–47^. We suspect the dual role may be associated with the syncytial lifestyle of Ashbya compared to uninucleate budding yeast. Future studies will be needed to mechanistically link the observed spatial translation variations across the cell cycle to generating asynchrony. Irrespective of the mechanism of Cln3-translation’s role in asynchrony, we propose that local variation in Cln3 translation emerges out of dynamic switching between a repressive and translation-permissive Whi3 states associated with the cell cycle.

Bni1 protein is enriched at hyphal tips^31^, and *BNI1* also shows a spatial bias in translation at tips but production appears infrequent as translation is predominantly repressed at hyphal tips. However, when Bni1 is being translated, it was associated with Whi3 signals. Taking live cell growth and condensate size observations together, we propose that Whi3 condensates associated with slower hyphal growth may serve to pulse translation at this site such that slow growing tips are translating Bni1 that then restore faster growth, concomitant with shrinking of Whi3 condensates. Notably, this allows *BNI1* transcripts to be positioned, poised for translation at the tip, but otherwise repressed, consistent with a low but essential demand for fresh protein at these sites. This repression is sensitive to signaling, and the Whi3 PKA phosphomimic, whi3(S637D) dramatically reverses repression both in vitro and in vivo. However, translation-state switching is also important for *BNI1*: this constitutively “active” mutant of Whi3 that allows high frequency of tip-associated translation results in abnormal overall cell morphology, with limited initiation of new polarity sites through lateral branching^5^. We hypothesize that the metered translation of Bni1 at tips is needed for overall balanced supply and production of Bni1 that enables many polarity sites to coexist to produce the normal branched morphology of the organism. Failure to form condensates results in abnormal growth phenotypes, including altered nuclear density, nuclear division synchrony, irregular branching interval and premature tip-splitting, possibly from unregulated excess BNI1 activity at existing sites ^3–5^. Notably, we rarely see these defects in WT cells, suggesting that nuclear division phenotypes are normally tightly co-regulated with branching and hyphal growth. This raises the intriguing possibility that by locally regulating a metered Cln3 and Bni1 abundance, Whi3 may serve the role of a coupler that is able to co-regulate nuclear division and hyphal growth in *Ashbya*’s shared cytoplasm to maintain coordinated growth.

Whi3 condensates, like many native condensates, hover around the diffraction limit, making them difficult to analyze at the subcondensate scale in vivo but the in vitro system brought some insights into how condensates may dually activate or repress translation. One of the most surprising findings in vitro is the specificity that the RNAs themselves bring to these condensates: *CLN3* 5’UTR shows consistent repression that is reversed by Whi3 phosphostate and binding site presence; *BNI1* shows more a complex biphasic response to Whi3 concentration that may be explained by differential dense-phase translation efficiencies. While we tested the contribution of Whi3 binding site abundance to translation regulation, we predict other factors such as RNA length, structure, motif position and clustering and 5’UTR features may all influence this specificity. The strongest RNA-independent predictor of translation outcome appears to be condensate size. Across all the Whi3 and RNA mutants and over a range of protein/RNA concentrations we test, the magnitude of bulk translation repression increases with condensate size, providing a compelling rationale for the tight size control seen in vivo^29^. Intriguingly, we also see condensate size and number variation in vivo that is associated with translation phenotypes: the largest Whi3 condensates appear at hyphal tips and are correlated with slower growth and variable *BNI1* translation at these sites, whereas M-phase nuclei are associated with both fewer, smaller condensates and highest *CLN3* translation. Thus size-dependent condensate function regulation may be a critical feature of condensate biology, as attested by the diverse biological systems in which it is starting to be reported^43,48^.

At the subcondensate scale that can be probed in vitro, we observe a striking translation-permissive zone in Whi3 condensates that is biased towards the condensate interface under specific compositions. The radial position of this zone is tunable by the valency of the RNA and Whi3 state. How condensate size and interfacial properties influence translation remains unknown. We observe a gradient of ribosome concentration across the condensates in vitro (**Figure 7E,F**), suggesting that the mesh-size of the condensate may in part limit accessibility of translation machinery depending on location in the dense phase and the condensate-enhanced translation effect at the interface may be coupled to enhanced concentration of translation machinery, as has been seen for other reactions in condensates^43^. Unfortunately, the RCA-PLA method does not allow us to quantify either the relative magnitude of translation or resolve spatial localization of translation within the condensate below the size of the amplicons in cells. Thus, the role of the condensate interface in promoting translation in vivo remains below the detection limit in this present study. The in vitro data presented here, however, suggest that the condensate surface may be an important zone to promote translation.

Another regime where bulk translation is repressed in vitro is in the dilute phase, where *CLN3* with WT Whi3 is modestly repressed. We propose that these repressive regimes are critical for the overall regulatory circuit: both Cln3 and Bni1 proteins are likely to be potent at low concentrations, when dosed in at the appropriate subcellular location. Consequently, bulk translation levels measured in vitro may not be the most relevant parameter in vivo: for instance, large condensates show the highest repression at the bulk level, but also show varying levels of interface-enriched translation, and these may be sufficient for shaping local Bni1 activity at the tip. In contrast, repression of Cln3 in the dilute phase may help accentuate local bursts of protein synthesis at certain cell-cycle states, allowing for enhanced spatial heterogeneity in composition.

This work highlights how condensates can function to toggle between distinct states to provide a dosed response, in this case to demands for specific proteins, in specific locations. The high degree of modulation of material states by changing binding interaction strengths and the ability of cells to control where and when condensates form make them powerful regulatory switches for local biochemistry.

## Author Contributions

A.S.G., Z.M.G., A.P.J., designed the study, interpreted results and wrote the initial version of the manuscript. A.P.J, S.J.C performed and analyzed *in vivo* growth measurements in *Ashbya*. A.P.J designed, performed, and analyzed in vivo spatial measurements of translation and ribosome profiling. Z.M.G. designed, performed, and analyzed *in vitro* translation assays. Z.M.G. designed and performed condensate purification for mass-spectrometry. Z.M.G., A.P.J., performed protein purification. All the authors read and approved the final manuscript.

## Acknowledgements

We thank Qiang Chen and Christopher Nicchitta for generous help with designing the polysome profiling experiments. We thank Christine Roden and Wilton Snead for their generous gift of N-protein and Fus, respectively. We thank Laura Herring and her team at the Michael Hooker Metabolomics and Proteomics Core located at the University of North Carolina at Chapel Hill. We thank Madeline Keenen, Veronica Farmer and Ben Stormo for providing useful comments on the manuscript.

## Funding

This work was supported Air Force Office of Scientific Research FA9550-20-1-0241, NIH grant 7R01GM081506-13, and 1R35GM156800 to ASG, Duke School of Medicine International Chancellor’s Scholarship, NIH F32 1F32GM147989 to APJ and F32 GM151858-02 to ZMG.

## Competing interests

The authors declare no competing interests.

## Inclusion & Ethics

This study did not include local researchers and did not impact the local community in any adverse ways.

## Code availability

No custom code central to the research was generated for this study.

## Data and materials availability

Data and materials are available upon request.

---

**Figure S1.**
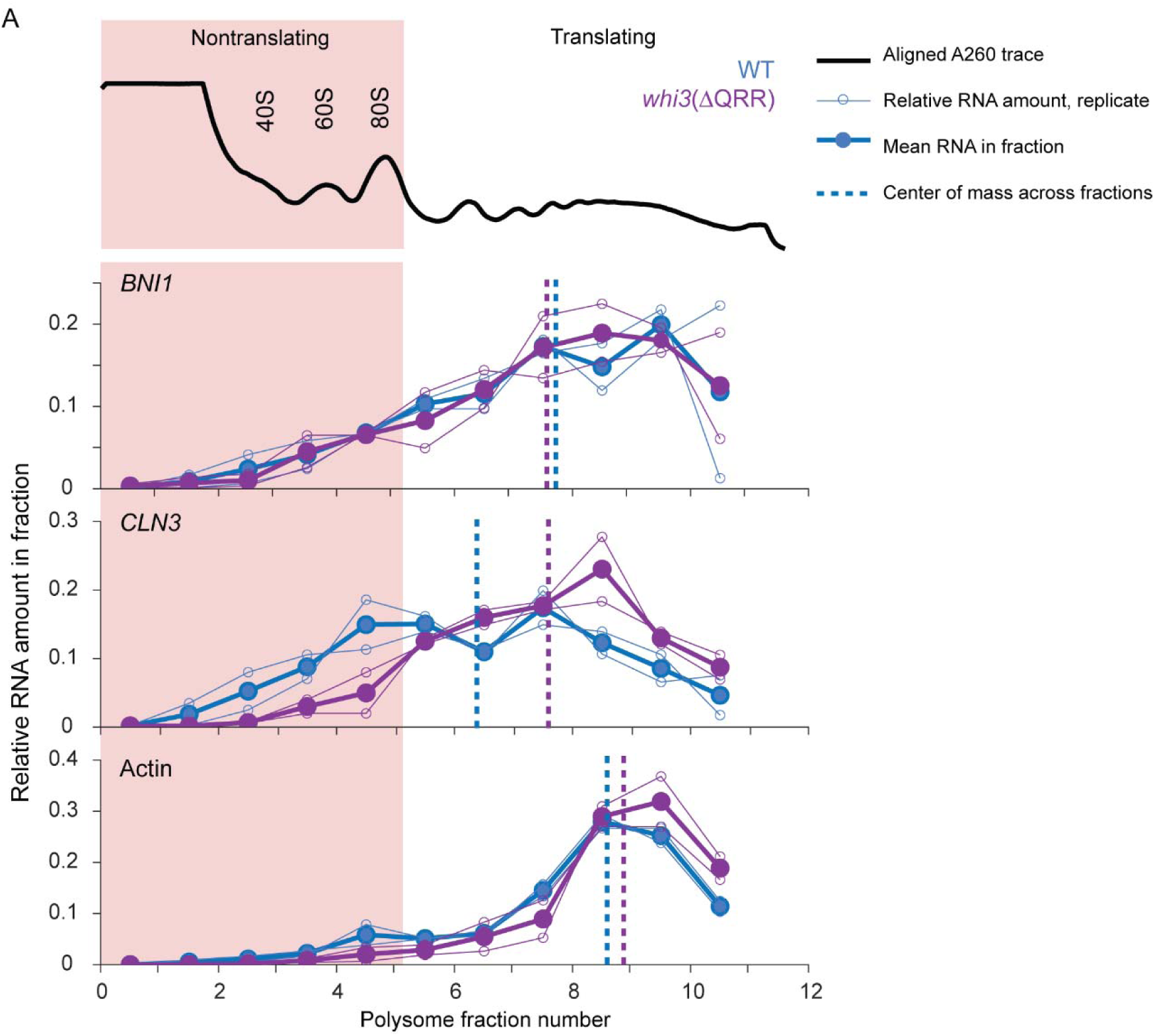
related to Figure 2. Whi3 condensates show enrichment of translation regulators. A. Polysome profile of *CLN3* and *BNI1* RNA levels in WT (blue) and deltaQRR (purple) indicates that these RNAs are polysome associated in WT, and *CLN3* shows a loss of a non-translating pool in *whi3*(ΔQRR), n= 2 independent sucrose gradients, whereas *BNI1* and *Actin* mRNAs don’t show significant shifts. Black line: representative A260 trace aligned to fractions; thin lines, open circles: individual replicates showing relative RNA abundance across fractions; thick lines, filled circles: average relative RNA of two replicates; dotted lines is the center-of-mass of the averaged RNA trace.

**Figure S2.**
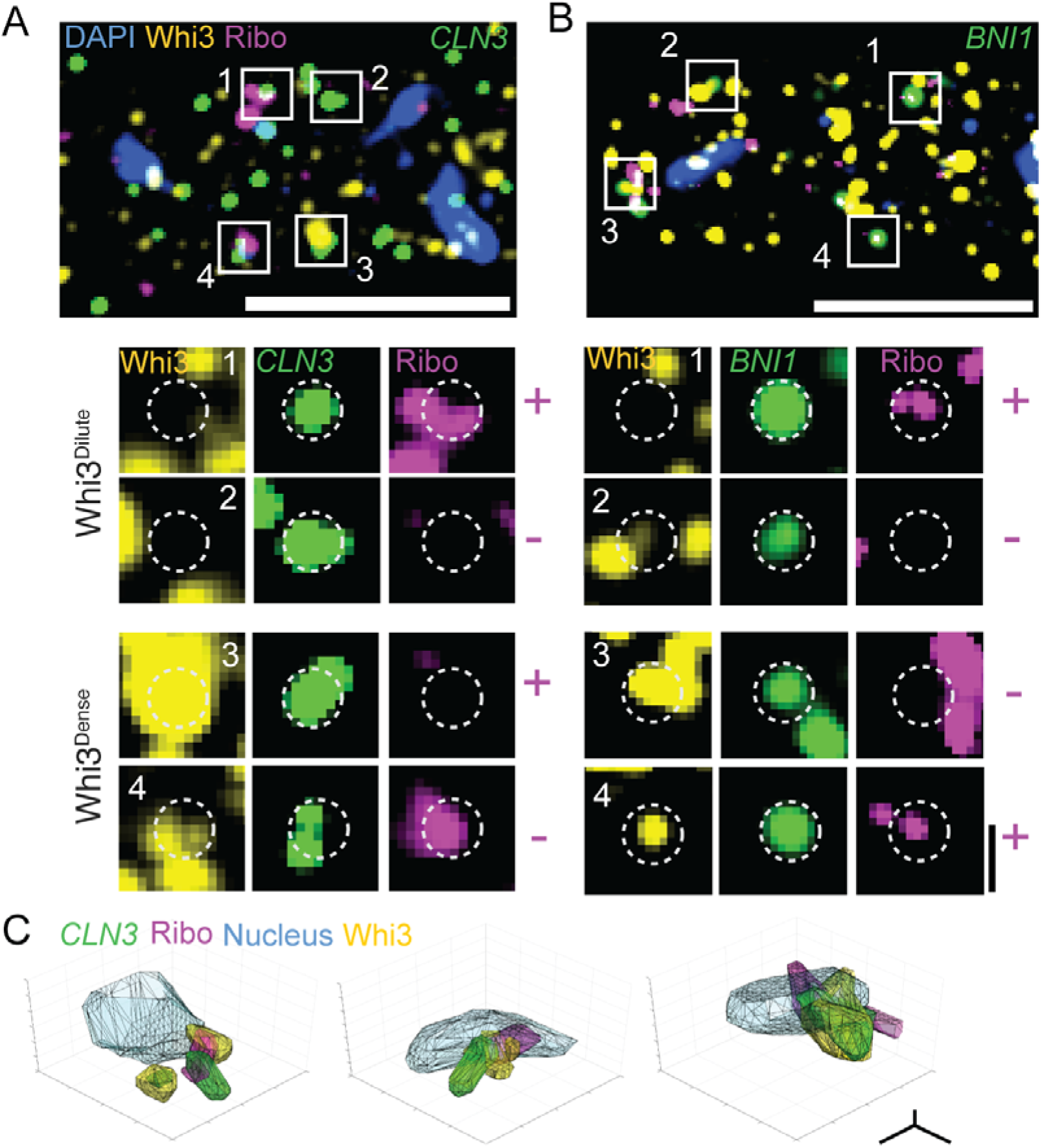
related to figure 4. Whi3 targets show variable translation at Whi3 dense and dilute phases. A. Representative image with insets showing examples of Whi3 dilute (1,2)- and dense-phase (3,4) associated *CLN3* RNAs showing dilute phase-associated translating (1), non translating (2) and dense-phase associated non-translating (3) and translating (4) RNA spots. Translation states is marked next to the insets; + = translation, - = non-translating. Scale bar 0.5 µm. B. Same as A for *BNI1*. C. Representative 3D reconstructions highlighting nuclear periphery associated Whi3 condensates showing translating *CLN3*, related to Figure 4A,C. Scale bar 0.5 microns.

**Figure S3.**
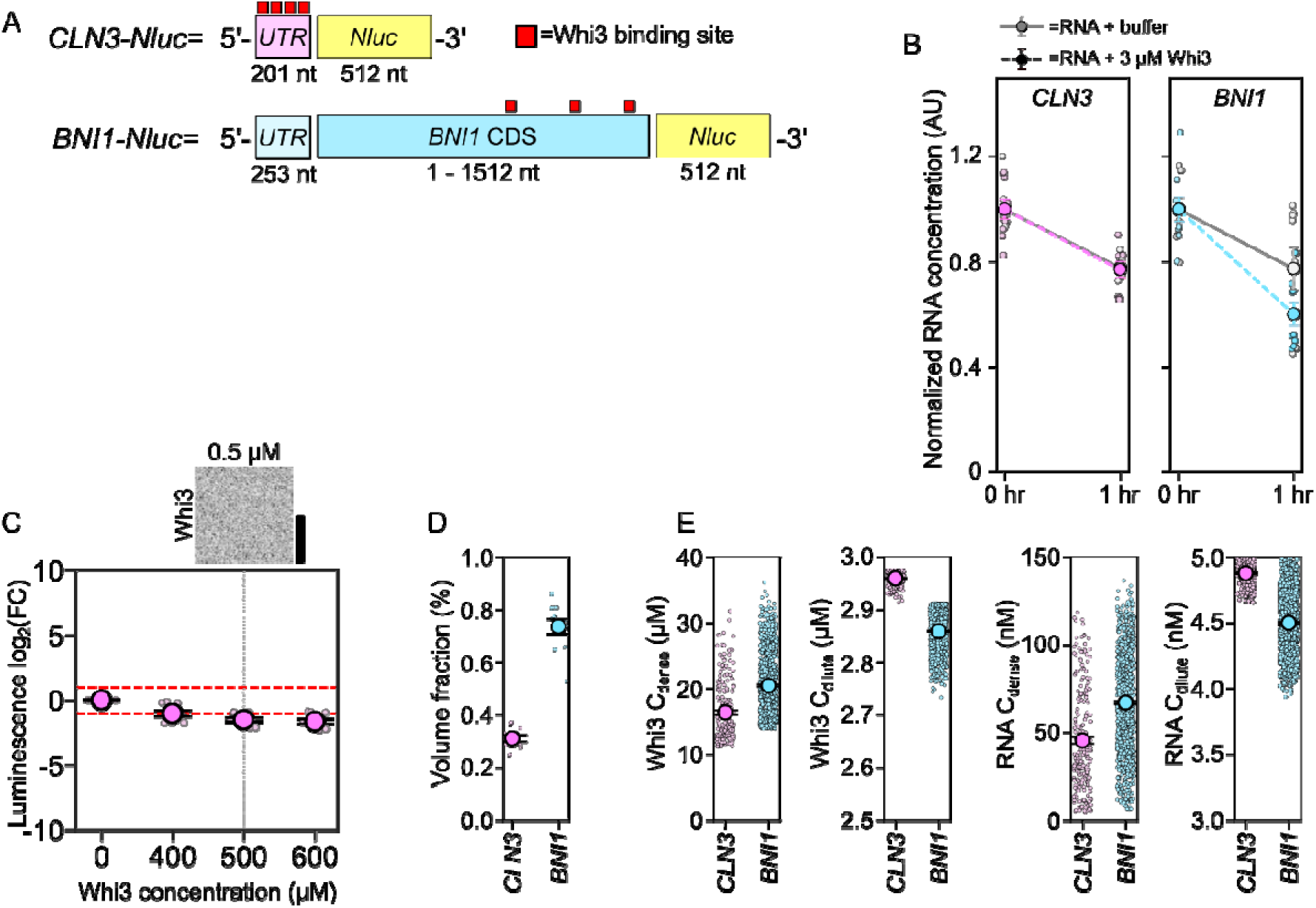
related to figure 5. Translation of *CLN3-Nluc* mRNA is repressed even in the absence of Whi3 condensates. A. Schematic of *CLN3-Nluc* and *BNI1-Nluc* reporter mRNAs. B. Normalized RNA concentration measured in RR lysates after the addition of 5 nM RNA (*BNI1* = lightblue, n=9; *CLN3* = pink, n=9) and 3 µM Whi3 and after one hour of translation. Reactions without Whi3 are shown in light grey. RNA concentrations are normalized to the average translation measured for that condition at time = 0. Measurements are taken from three separate RR lysates with three technical replicates per lysate. Means are displayed +/- SEM. C. Translation of *CLN3-Nluc* reporter RNA as a function of Whi3 concentration. Luminescence values are displayed as log_2_(FC) over RNA-only conditions. Measurements are taken from at least three separate RR lysates with three technical replicates per lysate (see Methods). Fluorescence image shows Whi3 signal at 0.5 µM. Scale bar = 2.5 µm. Means are displayed +/- SEM. D. Estimates of the dense phase volume fraction in Whi3 condensate reactions containing 3 µM Whi3 and either 5 nM *BNI1-Nluc* (light-blue; n=11) or CLN3-Nluc (pink; n=10). Each dot represents an imaging FOV. At least three FOVs were acquired per lysate. Means are displayed +/- SEM. E. Estimates of the dense and dilute phase concentrations for reactions containing 3 µM Whi3 and either 5 nM BNI1-Nluc (light-blue; n=896) or CLN3-Nluc (pink; n=224). Means are displayed +/- SEM.

**Figure S4.**
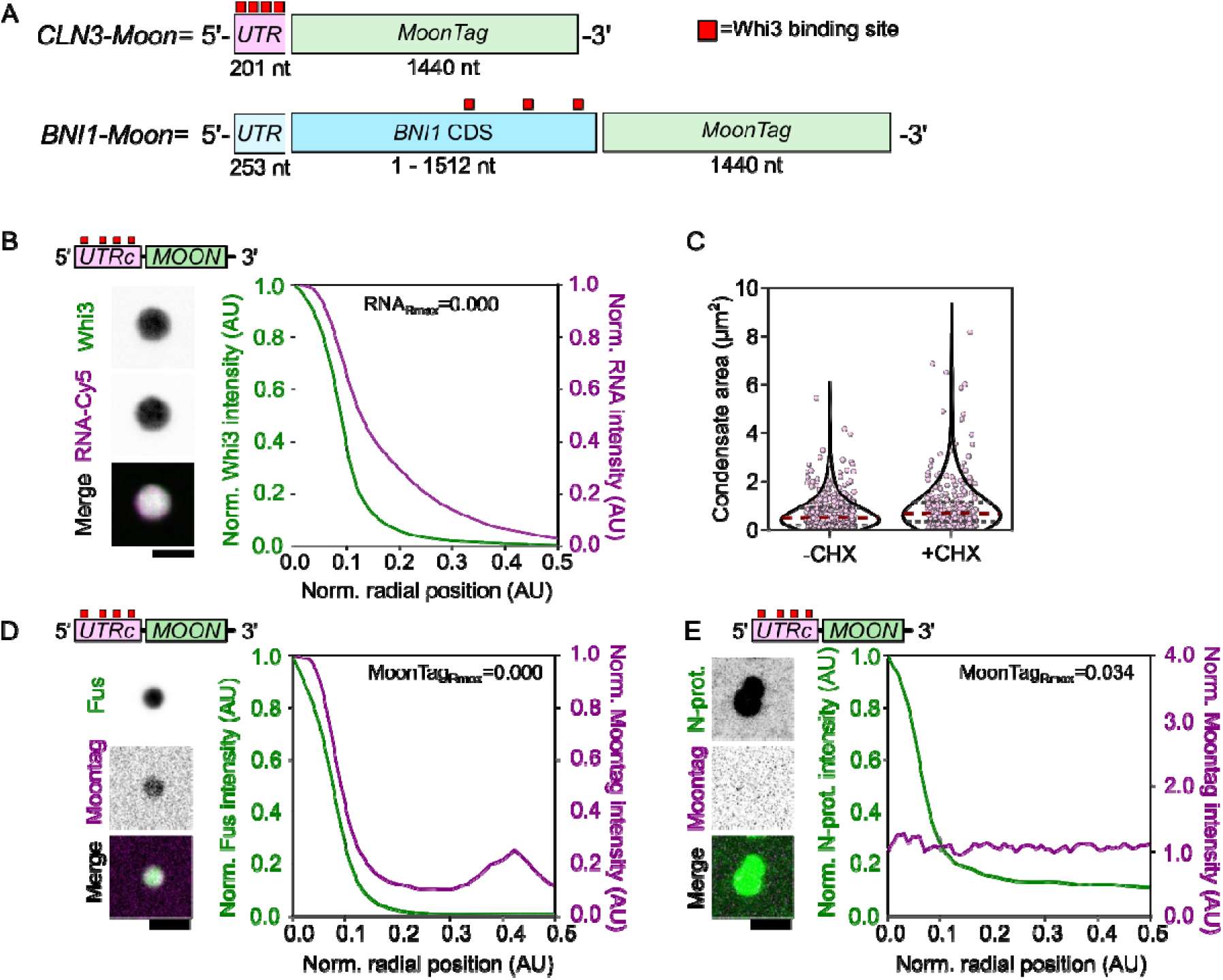
related to figure 6. mRNA valency influences Whi3 condensate translation. A. Schematic showing *BNI1-Moon* and *CLN3-Moon* reporter mRNAs. B. Fluorescence images of Whi3 condensates formed with *Cy5-CLN3-Moon*. Radial intensity profile of Whi3 condensates formed with *Cy5-CLN3-Moon*, showing fluorescence intensity as a function of radial position in the condensate for Whi3 and Cy5-RNA. 0.1 AU from the center of the condensate is equal to the radius of the condensate (dotted grey line). The radial position associated with the maximum RNA intensity recorded, RNA_Rmax_, is shown. Trace shows the average intensity of 55 condensates ± SEM. Scale bar = 2.5 µm. C. Whi3 condensate area for *CLN3-Moon* condensates in the translation competent RR lysates (CHX-; 435) or translation inhibited RR lysates (CHX+; n=255). Individual dots indicate a single condensate. D. Fluorescence images of Fus condensates formed with *CLN3-Moon* showing accumulation of Nb-mCherry within the Fus condensate in cycloheximide treated extracts. Radial intensity profile of Fus condensates formed with *CLN3-Moon*, showing fluorescence intensity as a function of radial position in the condensate for Fus or Nb-mCherry. Scale bar = 2.5 µm. 0.1 AU from the center of the condensate is equal to the radius of the condensate (dotted grey line). The radial position associated with the maximum MoonTag intensity recorded, MoonTagRmax, is shown. Trace shows the average intensity of 265 condensates ± SEM. E. Fluorescence images of N-protein condensates formed with *CLN3-Moon* showing no translation of mRNAs or accumulation of Nb-mCherry within or at the surface of the N-protein condensate. Radial intensity profile of N-protein condensates formed with *CLN3-Moon*, showing fluorescence intensity as a function of radial position in the condensate for N-protein or MoonTag Nb-mCherry. 0.1 AU from the center of the condensate is equal to the radius of the condensate (dotted grey line). The radial position associated with the maximum MoonTag intensity recorded, MoonTagRmax, is shown. Trace shows the average intensity of 162 condensates ± SEM. Scale bar = 2.5 µm.

**Figure S5.**
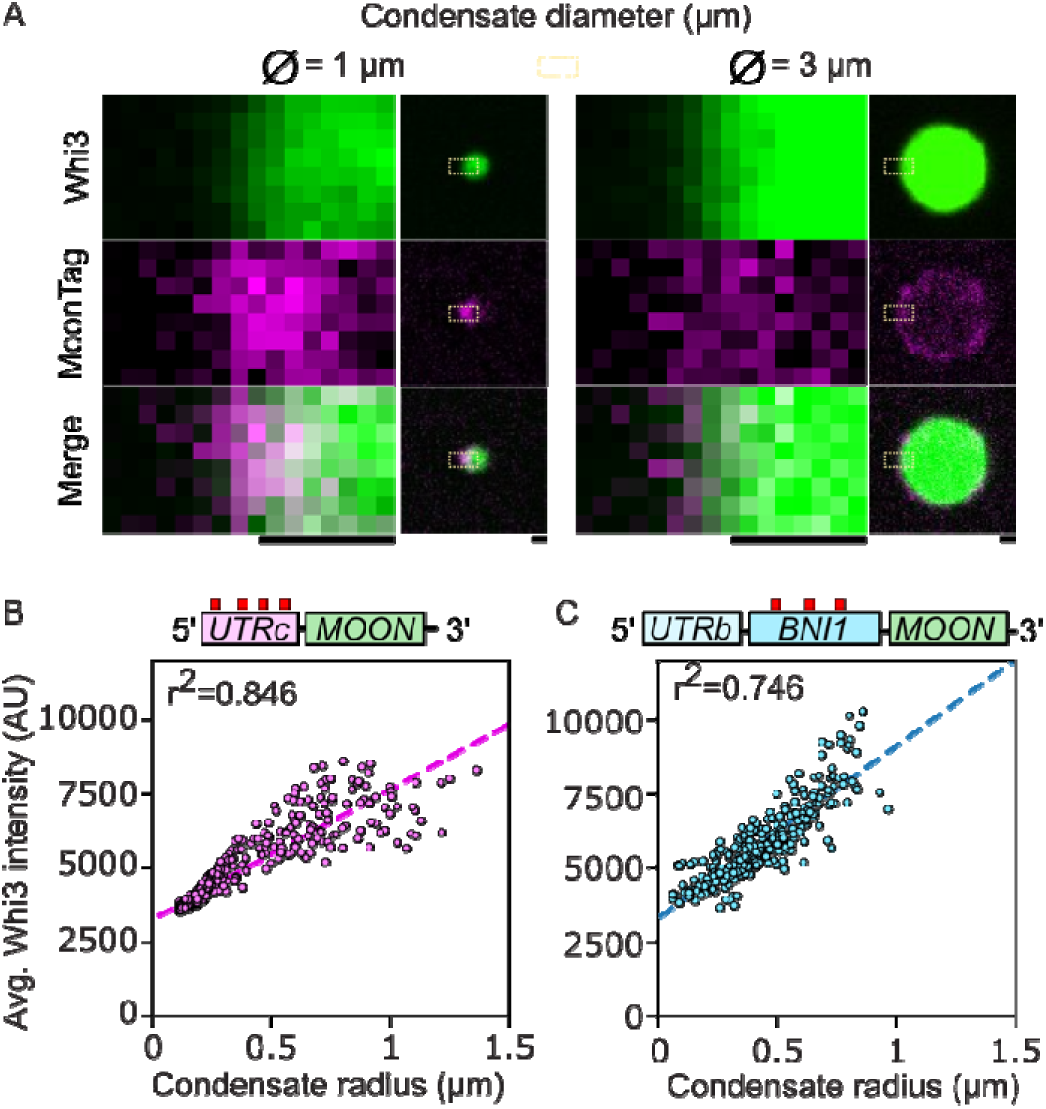
related to figure 6. Whi3 concentration scales with condensate size. A. 2D-fluorescence image representing the condensate interface for small (ø=1) and large (ø=3) condensates formed with *BNI1-Moon*. Scale bar = 0.5 µm. B. Whi3 concentration as a function of condensate radius for Whi3 condensates formed with *BNI1-Moon*. Each individual dot represents a condensate (n=300). C. Whi3 concentration as a function of condensate radius for Whi3 condensates formed with *CLN3-Moon* (n=854). Each individual dot represents a condensate.

**Figure S6.**
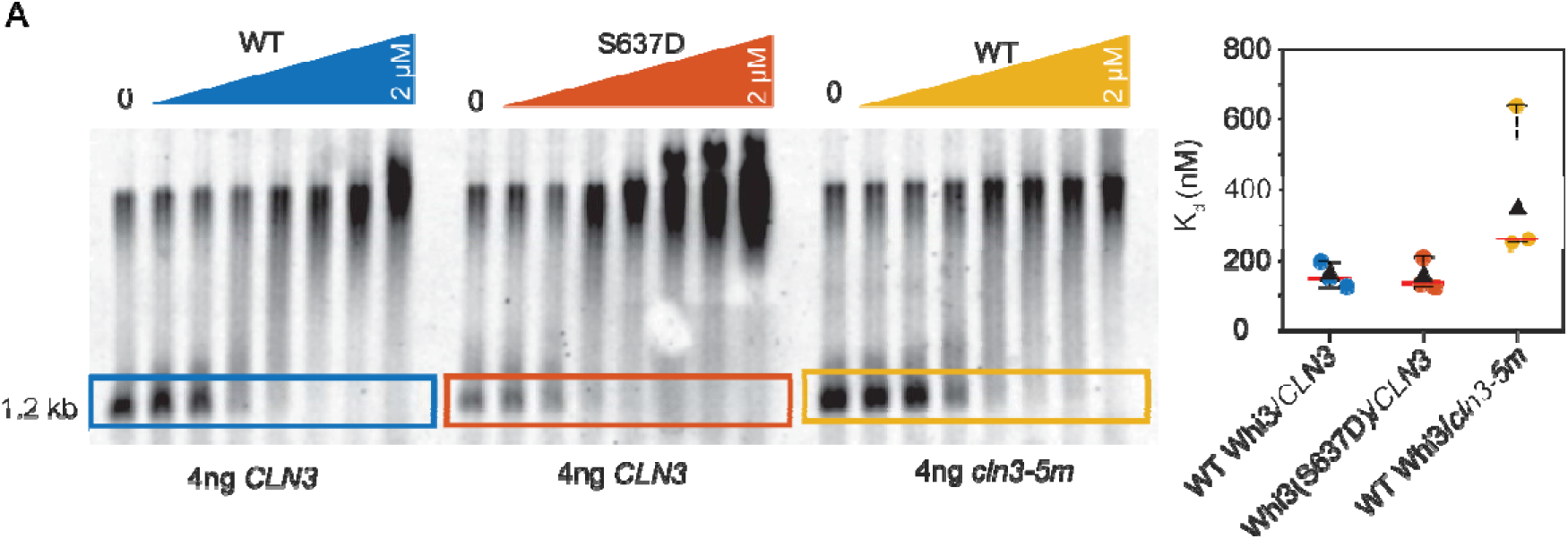
related to figure 5 and 6. Phospho-state impacts Whi3 target translation independently of binding site affinity. A. Apparent dissociation constants for WT or 0xWBS *CLN3* RNA with WT or S637D Whi3 from EMSAs. Three technical replicates of WTWhi3/WT cln3, S637D Whi3/WT CLN3 and WT Whi3/*cln3* 0xWBS shows that S637D does not severely change apparent Kd (both ∼150 nM), relative to WT, whereas WT Whi3 shows decreased affinity for the RNA variant with no binding sites.

## Notes

### Competing Interest Statement

The authors have declared no competing interest.

### Summary of Updates

Manuscript updated to include data supporting Whi3 condensates as sites of translation in vivo. The text has been updated to make more clear the functions and properties of dense and dilute phases. Additionally, descriptions of protein localizations where known, imaging of ribosomes within Whi3 condensates, measurements of RNA degradation in the presence of condensates in vitro, and normalization of MoonTag translation to condensate RNA concentrations have been added to the manuscript.

