## Supplementary material for "RNA-specific local translation is patterned by condensates for multinucleate cell growth": Methods

### Protein purification and labeling

Whi3, whi3(S637D), and whi3(ΔQRR) were purified and labeled with amine-reactive, NHS ester-functionalized Atto488 (Atto-Tec; AD 488) using the protocol previously reported in ^1^. The method was modified to exclude the 0.3 mg/mL lysozyme included during the cell lysis step. Prior to the addition of protein to experiments, all protein stocks were centrifuged at approximately 16,000 RCF for 5 min to remove large aggregates.

2H10-mCherry was expressed as an N-terminal 6his-fusion construct with a TEV cleavage site located after the N-terminal 6xHis purification tag. BL21 *E. coli* cells were induced with 1 mM IPTG for 16 h at 18 °C. Cells were harvested by centrifugation, and bacterial pellets were resuspended in 25 mL lysis buffer (20 mM HEPES (pH 7.4), 100 mM KCl, and 1x Pierce Protease inhibitor tablet (A32955)). Lysis buffers containing the resuspended bacterial pellets were then transferred to a Dounce-homogenizer before probe sonication. Protein was purified from crude bacterial extract by incubating 25 mL of bacterial lysate with 0.8 mL of cobalt resin slurry (0.4 mL packed beads). After 1h of incubation with the cobalt slurry, bacterial lysates were loaded onto gravity column and allowed to settle (~20 min). The gravity column was then extensively washed using excess lysis buffer (at least 10x column volumes). Protein was eluted in 3 mL of elution buffer (200 mM imidazole, 20 mM Hepes pH 7.4, 100 mM KCl) added in 1mL increments. Protein was dialyzed overnight against the same buffer without imidazole. Protein concentration was determined by Bradford. Protein was flash-frozen and stored as small aliquots at −80 °C. Prior to the addition of 2H10-mCherry to translation reactions, purified 2H10-mCherry was mixed with TEV protease in a 17:1 ratio (2H10 to TEV) and allowed to incubate for ~20 min at RT which resulted in sufficient TEV cleavage and subsequently 2H10 functional recognition of MoonTag peptides. Prior to the addition of 2H10-mCherry to experiments, the aliquot was centrifuged at approximately 16,000 rcf for 5 min to remove large aggregates.

### Cloning

Full-length Nanoluciferase (Nluc; Addgene, plasmid #60140) was amplified using primers containing overlap with the 3′ end of either the *CLN3* 5′ UTR, or the CDS of *BNI1*. The following Nluc fragments containing overlap with *CLN3* or *BNI1* respectively, were included in a PCR reaction containing the respective fragments of *CLN3* or *BNI1*, allowing for the amplification of the *CLN3-Nluc* and *BNI1-Nluc*. After the amplification of *BNI1-Nluc*, an additional round of PCR was carried out using primers containing overlap to the 5′UTR of *BNI1*. *BNI1-Nluc* (containing overlap with *BNI1*’s 5′UTR) and *BNI1*’s 5′UTR were included in a PCR reaction allowing for the amplification of *UTR-BNI1-Nluc*. PCR products were then cloned into the pJet vector (Thermofisher Scientific K1231) using blunt end cloning.

A custom MoonTag sequence lacking any UGCAUs or GCAUs (Supplemental Table 1) was ordered and subsequently cloned into the pJet vector (Thermofisher Scientific K1231) using blunt end cloning. To insert the 5′ UTR of *CLN3* or the CDS of *BNI1* into the custom MoonTag cassette, *CLN3* or *BNI1* gene fragments were amplified using primers containing a 5′ DraIII and 3′ BsiWI restriction sites and cloned into the MoonTag cassette using restriction enzyme digest of both insert and vector.

| Sequence ID | Sequence |
| --- | --- |
| *CLN3-Nluc* | TAATACGACTCACTATAAGGGTCTGCATACCAAGGATCAGCCGCTTGCATTAAAGGGGACGAACCGGGGCTCTTATCCTTAGCATCGTCTCTTGTCCACACCTCGTTGACCTGCAGACACAGCACAAACCCTGCATAATTAATAGTAATCATTGTCATCTCCAGCTGGGCTGTTAATATCTCATACCCGAACACCTTTGCATATTATACACATATTCGTCATGGTCTTCACACTCGAAGATTTCGTTGGGGACTGGCGACAGACAGCCGGCTACAACCTGGACCAAGTCCTTGAACAGGGAGGTGTGTCCAGTTTGTTTCAGAATCTCGGGGTGTCCGTAACTCCGATCCAAAGGATTGTCCTGAGCGGTGAAAATGGGCTGAAGATCGACATCCATGTCATCATCCCGTATGAAGGTCTGAGCGGCGACCAAATGGGCCAGATCGAAAAAATTTTTAAGGTGGTGTACCCTGTGGATGATCATCACTTTAAGGTGATCCTGCACTATGGCACACTGGTAATCGACGGGGTTACGCCGAACATGATCGACTATTTCGGACGGCCGTATGAAGGCATCGCCGTGTTCGACGGCAAAAAGATCACTGTAACAGGGACCCTGTGGAACGGCAACAAAATTATCGACGAGCGCCTGATCAACCCCGACGGCTCCCTGCTGTTCCGAGTAACCATCAACGGAGTGACCGGCTGGCGGCTGTGCGAACGCATTCTGGCGGGTGGGGGATCATTGTACAAATAA |
| *cln3(4m)-Nluc* | TAATACGACTCACTATAAGGGACGTCCTAGTACCAAGGATCAGCCGCTCTAGTTAAAGGGGACGAACCGGGGCTCTTATCCTTAGCATCGTCTCTTGTCCACACCTCGTTGACCTGCAGACACAGCACAAACCCCTAGTAATTAATAGTAATCATTGTCATCTCCAGCTGGGCTGTTAATATCTCATACCCGAACACCTTCTAGTATTATACACATATTCGTCATGGTCTTCACACTCGAAGATTTCGTTGGGGACTGGCGACAGACAGCCGGCTACAACCTGGACCAAGTCCTTGAACAGGGAGGTGTGTCCAGTTTGTTTCAGAATCTCGGGGTGTCCGTAACTCCGATCCAAAGGATTGTCCTGAGCGGTGAAAATGGGCTGAAGATCGACATCCATGTCATCATCCCGTATGAAGGTCTGAGCGGCGACCAAATGGGCCAGATCGAAAAAATTTTTAAGGTGGTGTACCCTGTGGATGATCATCACTTTAAGGTGATCCTGCACTATGGCACACTGGTAATCGACGGGGTTACGCCGAACATGATCGACTATTTCGGACGGCCGTATGAAGGCATCGCCGTGTTCGACGGCAAAAAGATCACTGTAACAGGGACCCTGTGGAACGGCAACAAAATTATCGACGAGCGCCTGATCAACCCCGACGGCTCCCTGCTGTTCCGAGTAACCATCAACGGAGTGACCGGCTGGCGGCTGTGCGAACGCATTCTGGCGGGTGGGGGATCATTGTACAAATAA |
| *BNI1-Nluc* | GAAAATTGTCCCAATTAGTAGCATCACGCTTAATACGACTCACTATAAGACTAACATATAGAGGGCAGTACAGCACACGTCAACTATCTTTGCTCGTCGATCGTCGGGCATTTACACACCACAGTACACGCGCGCGCCTGGGGCTTGGAGATACATTTCTGCTTGCAGGCCCGAATATTGCGCTAATCGTCAAGGCACTCCAGCATTCCACGCAGTGGGACAAGGCAGCAGGTGTGGAACACGGCGGGACGCAGGATTTACCGGTATAGTGGCAGCGTCGGCGGCTGGGCACATATGCAAGGGCCACCATGAAGAAGTCCACGCACTCGAACAAACACTCCAAAACACAGCACGCGGGGCAGGTTGCCGGGGCAACACACGCCAACGGTGGCCACAGCGGCGGTGGCTTGTTCTCGAACCTCATGAAGTTGACGGGCTCGTCGGGCTCGCAGAGGATTGCGACCTCGGATATTTCGTCTCCGAAGAAGGTGACGCTGCCGTCCAATCTGGACCTGTCCGGTGCAAAACCCTTGAACAAGCTGAATACGATCAACGCGTCCAATTTACCACAGGTGTCGGGCGAGGTAACACAGGCCCGATCTGCGTCTGGCGCTGCGTTGTCTTCACCATCCAAGTATTCATATTCCAAGCGGTCTTCCCAGTGGGCGAGCTCCAAGCAAGCAAACCATCCAGAACACGACCAGCAGTATCCCCAGCATCTTAACCATCTTTTTGTGCAGCCCCATCCTCACCACTCGGCTTTGACGCGGCAGATGACCAACCAATCAAATTACTCCGCTTCTTCATATACTTCTTTATCTCTGCTACATAAAGCCACGGATATAGATGGTACCTTAACCTTGGAGAAGCCCGAGGATCCCCAAGAGATTGAAGAACTATACCAGGAGTTGCTACAGAAAAGAAACATACCACAATCCGTTTCTGTCCACGGTCATCGCGAGCTGATGAGTTATGGCATTGATAAAAAGTGGCTTATGGTTAAACAGGATCTCCAAACGGAATACAAGAAGATGAAAAATAGCATGCCACCAAGCTCGACCGTAGAGTCGATAGCGAGTTCTGCTCTAGCAAAATCCCCTGGCGGAAGCATATCGTCCGGGAGCATCACAAGATCCATATCTCAAAACAGCGCGAAGCCCATGGCTGTAGCCTACGAGCCTGCACATCTCTCGCCCGATCACTACGTGCGGAAGATTATATCGGATAGAGTGACTGCACAAGAACTAAACGATTTGTGGATTTCGTTGAGGACCGAAAGCATAGACTGGGTGATTGGTTTTATCGACGCTCAGGGACAGGTTGCAATTGCCAATCGCCTTTTAAAATTTGTGCAACGTGAGTCTCTAGATATTCTACATGATTATGATGCATTAGAGAAGGAAAACGCCTACTATAAGTGTTTGAGAGTCCTTACGAACTTGCGGGAAGGGATGCAGGAAGCGCTGAAGTCCAAACTAGTGATCAGCTCTGTGGTGGAGGGTCTATTATCCAGTAGGTTGTCCACCAGGAGGGTAGCAACGGAGACTCTATTGTACATGCTCTCTGACGACGAGCTCAAAACTGCAGATACGGGTCAAGATGCGATTTTTCCTTTGTTGGTTGCATTAGACCAAGAATCAAAGTTTGCTGCAAACATTCATCTTCGTGGAAGACTTCATGAAACTAATAAACGTAAATCCTTACCTACTGGGAGAGCAGATTTCCATGTGGTCGCCAAACGACTAGAACAATGGCTATATGTCATTGAATATACCTTGGACGGTCGCGGGAGAATGGGTTCTCTTGTTGGTGCATCTGAGGTCTTCACACTCGAAGATTTCGTTGGGGACTGGCGACAGACAGCCGGCTACAACCTGGACCAAGTCCTTGAACAGGGAGGTGTGTCCAGTTTGTTTCAGAATCTCGGGGTGTCCGTAACTCCGATCCAAAGGATTGTCCTGAGCGGTGAAAATGGGCTGAAGATCGACATCCATGTCATCATCCCGTATGAAGGTCTGAGCGGCGACCAAATGGGCCAGATCGAAAAAATTTTTAAGGTGGTGTACCCTGTGGATGATCATCACTTTAAGGTGATCCTGCACTATGGCACACTGGTAATCGACGGGGTTACGCCGAACATGATCGACTATTTCGGACGGCCGTATGAAGGCATCGCCGTGTTCGACGGCAAAAAGATCACTGTAACAGGGACCCTGTGGAACGGCAACAAAATTATCGACGAGCGCCTGATCAACCCCGACGGCTCCCTGCTGTTCCGAGTAACCATCAACGGAGTGACCGGCTGGCGGCTGTGCGAACGCATTCTGGCGGGTGGGGGATCATTGTACAAATAA |
| *bni1(3m)-Nluc* | GAAAATTGTCCCAATTAGTAGCATCACGCTTAATACGACTCACTATAAGACTAACATATAGAGGGCAGTACAGCACACGTCAACTATCTTTGCTCGTCGATCGTCGGGCATTTACACACCACAGTACACGCGCGCGCCTGGGGCTTGGAGATACATTTCTGCTTGCAGGCCCGAATATTGCGCTAATCGTCAAGGCACTCCAGCATTCCACGCAGTGGGACAAGGCAGCAGGTGTGGAACACGGCGGGACGCAGGATTTACCGGTATAGTGGCAGCGTCGGCGGCTGGGCACATATGCAAGGGCCACCATGAAGAAGTCCACGCACTCGAACAAACACTCCAAAACACAGCACGCGGGGCAGGTTGCCGGGGCAACACACGCCAACGGTGGCCACAGCGGCGGTGGCTTGTTCTCGAACCTCATGAAGTTGACGGGCTCGTCGGGCTCGCAGAGGATTGCGACCTCGGATATTTCGTCTCCGAAGAAGGTGACGCTGCCGTCCAATCTGGACCTGTCCGGTGCAAAACCCTTGAACAAGCTGAATACGATCAACGCGTCCAATTTACCACAGGTGTCGGGCGAGGTAACACAGGCCCGATCTGCGTCTGGCGCTGCGTTGTCTTCACCATCCAAGTATTCATATTCCAAGCGGTCTTCCCAGTGGGCGAGCTCCAAGCAAGCAAACCATCCAGAACACGACCAGCAGTATCCCCAGCATCTTAACCATCTTTTTGTGCAGCCCCATCCTCACCACTCGGCTTTGACGCGGCAGATGACCAACCAATCAAATTACTCCGCTTCTTCATATACTTCTTTATCTCTGCTACATAAAGCCACGGATATAGATGGTACCTTAACCTTGGAGAAGCCCGAGGATCCCCAAGAGATTGAAGAACTATACCAGGAGTTGCTACAGAAAAGAAACATACCACAATCCGTTTCTGTCCACGGTCATCGCGAGCTGATGAGTTATGGCATTGATAAAAAGTGGCTTATGGTTAAACAGGATCTCCAAACGGAATACAAGAAGATGAAAAATAGCATGCCACCAAGCTCGACCGTAGAGTCGATAGCGAGTTCTGCTCTAGCAAAATCCCCTGGCGGAAGCATATCGTCCGGGAGCATCACAAGATCCATATCTCAAAACAGCGCGAAGCCCATGGCTGTAGCCTACGAGCCTGCACATCTCTCGCCCGATCACTACGTGCGGAAGATTATATCGGATAGAGTGACTGCACAAGAACTAAACGATTTGTGGATTTCGTTGAGGACCGAAAGCATAGACTGGGTGATTGGTTTTATCGACGCTCAGGGACAGGTTGCAATTGCCAATCGCCTTTTAAAATTTGTGCAACGTGAGTCTCTAGATATTCTACATGATTATGACGCTTTAGAGAAGGAAAACGCCTACTATAAGTGTTTGAGAGTCCTTACGAACTTGCGGGAAGGGATGCAGGAAGCGCTGAAGTCCAAACTAGTGATCAGCTCTGTGGTGGAGGGTCTATTATCCAGTAGGTTGTCCACCAGGAGGGTAGCAACGGAGACTCTATTGTACATGCTCTCTGACGACGAGCTCAAAACTGCAGATACGGGTCAAGATGCGATTTTTCCTTTGTTGGTAGCCTTAGACCAAGAATCAAAGTTTGCTGCAAACATTCATCTTCGTGGAAGACTTCATGAAACTAATAAACGTAAATCCTTACCTACTGGGAGAGCAGATTTCCATGTGGTCGCCAAACGACTAGAACAATGGCTATATGTCATTGAATATACCTTGGACGGTCGCGGGAGAATGGGTTCTCTTGTTGGGGCTTCTGAGGTCTTCACACTCGAAGATTTCGTTGGGGACTGGCGACAGACAGCCGGCTACAACCTGGACCAAGTCCTTGAACAGGGAGGTGTGTCCAGTTTGTTTCAGAATCTCGGGGTGTCCGTAACTCCGATCCAAAGGATTGTCCTGAGCGGTGAAAATGGGCTGAAGATCGACATCCATGTCATCATCCCGTATGAAGGTCTGAGCGGCGACCAAATGGGCCAGATCGAAAAAATTTTTAAGGTGGTGTACCCTGTGGATGATCATCACTTTAAGGTGATCCTGCACTATGGCACACTGGTAATCGACGGGGTTACGCCGAACATGATCGACTATTTCGGACGGCCGTATGAAGGCATCGCCGTGTTCGACGGCAAAAAGATCACTGTAACAGGGACCCTGTGGAACGGCAACAAAATTATCGACGAGCGCCTGATCAACCCCGACGGCTCCCTGCTGTTCCGAGTAACCATCAACGGAGTGACCGGCTGGCGGCTGTGCGAACGCATTCTGGCGGGTGGGGGATCATTGTACAAATAA |
| *Custom_MoonTag* | atgcgtacgggcaagaatgagcaggaactgctggaactggacaaatgggcaagcctggggagcggctccggaaagaacgagcaggaactgctggaactggacaagtgggcttctctgggctccgggtccgggaaaaacgaacaggagctgctggagctggataagtgggccagcctgggctccgggtccgggaagaatgaacaggaactgctggaactggataaatgggcctctctgggcagcgggtccggaaagaatgaacaggagctgctggaactggacaagtgggcgtccctggggagcgggtccggaaagaatgaacaggaactgctggaactggataagtgggcttccctgggaagcggctccggaaaaaacgagcaggagctgctggagctggacaagtgggcctctctgggctccggctccgggaagaatgaacaggagctgctggaactggataagtgggcctctctggggtccggaagcggtaaaaacgagcaggaactgctggaactggacaagtgggcaagcctggggtctggctccggcaaaaacgaacaggaactgctggaactggacaaatgggccagcctgggctccgggtccgggaaaaatgagcaggaactgctggaactggataaatgggcgagcctgggcagcgggtccggaaaaaacgaacaggagctgctggagctggacaaatgggcgagtctgggctccggaagcggcaagaatgaacaggaactgctggagctggacaaatgggccagcctgggcagcggctccggaaagaatgagcaggaactgctggagctggacaaatgggcgtctctgggctccggctccgggaaaaacgagcaggaactgctggagctggacaagtgggcttccctgggcagcggctccggaaaaaatgaacaggaactgctggaactggacaagtgggccagcctggggtccgggtctgggaagaacgagcaggaactgctggagctggacaagtgggcttctctggggtctggctctggcaagaacgagcaggaactgctggaactggacaagtgggcgagcctgggcagcgggtctggcaagaatgaacaggagctgctggaactggacaaatgggccagcctggggtctgggtctggaaagaatgagcaggaactgctggagctggacaagtgggcctctctgggctctgggtctggcaagaacgagcaggagctgctggagctggataagtgggcgtccctggggtccgggtccggcaagaatgaacaggagctgctggagctggataaatgggcttccctgggctctgggtctgggaagaatgagcaggagctgctggaactggataagtgggcgagcctgggcagcgggtctggcaagaatgagcaggaactgctggagctggacaagtgggcttctctg |
| *CLN3-MoonTag* | TAATACGACTCACTATAAGGGTCTGCATACCAAGGATCAGCCGCTTGCATTAAAGGGGACGAACCGGGGCTCTTATCCTTAGCATCGTCTCTTGTCCACACCTCGTTGACCTGCAGACACAGCACAAACCCTGCATAATTAATAGTAATCATTGTCATCTCCAGCTGGGCTGTTAATATCTCATACCCGAACACCTTTGCATATTATACACATATTCGTCGCCACCATGCGTACGGGCAAGAATGAGCAGGAACTGCTGGAACTGGACAAATGGGCAAGCCTGGGGAGCGGCTCCGGAAAGAACGAGCAGGAACTGCTGGAACTGGACAAGTGGGCTTCTCTGGGCTCCGGGTCCGGGAAAAACGAACAGGAGCTGCTGGAGCTGGATAAGTGGGCCAGCCTGGGCTCCGGGTCCGGGAAGAATGAACAGGAACTGCTGGAACTGGATAAATGGGCCTCTCTGGGCAGCGGGTCCGGAAAGAATGAACAGGAGCTGCTGGAACTGGACAAGTGGGCGTCCCTGGGGAGCGGGTCCGGAAAGAATGAACAGGAACTGCTGGAACTGGATAAGTGGGCTTCCCTGGGAAGCGGCTCCGGAAAAAACGAGCAGGAGCTGCTGGAGCTGGACAAGTGGGCCTCTCTGGGCTCCGGCTCCGGGAAGAATGAACAGGAGCTGCTGGAACTGGATAAGTGGGCCTCTCTGGGGTCCGGAAGCGGTAAAAACGAGCAGGAACTGCTGGAACTGGACAAGTGGGCAAGCCTGGGGTCTGGCTCCGGCAAAAACGAACAGGAACTGCTGGAACTGGACAAATGGGCCAGCCTGGGCTCCGGGTCCGGGAAAAATGAGCAGGAACTGCTGGAACTGGATAAATGGGCGAGCCTGGGCAGCGGGTCCGGAAAAAACGAACAGGAGCTGCTGGAGCTGGACAAATGGGCGAGTCTGGGCTCCGGAAGCGGCAAGAATGAACAGGAACTGCTGGAGCTGGACAAATGGGCCAGCCTGGGCAGCGGCTCCGGAAAGAATGAGCAGGAACTGCTGGAGCTGGACAAATGGGCGTCTCTGGGCTCCGGCTCCGGGAAAAACGAGCAGGAACTGCTGGAGCTGGACAAGTGGGCTTCCCTGGGCAGCGGCTCCGGAAAAAATGAACAGGAACTGCTGGAACTGGACAAGTGGGCCAGCCTGGGGTCCGGGTCTGGGAAGAACGAGCAGGAACTGCTGGAGCTGGACAAGTGGGCTTCTCTGGGGTCTGGCTCTGGCAAGAACGAGCAGGAACTGCTGGAACTGGACAAGTGGGCGAGCCTGGGCAGCGGGTCTGGCAAGAATGAACAGGAGCTGCTGGAACTGGACAAATGGGCCAGCCTGGGGTCTGGGTCTGGAAAGAATGAGCAGGAACTGCTGGAGCTGGACAAGTGGGCCTCTCTGGGCTCTGGGTCTGGCAAGAACGAGCAGGAGCTGCTGGAGCTGGATAAGTGGGCGTCCCTGGGGTCCGGGTCCGGCAAGAATGAACAGGAGCTGCTGGAGCTGGATAAATGGGCTTCCCTGGGCTCTGGGTCTGGGAAGAATGAGCAGGAGCTGCTGGAACTGGATAAGTGGGCGAGCCTGGGCAGCGGGTCTGGCAAGAATGAGCAGGAACTGCTGGAGCTGGACAAGTGGGCTTCTCTGTAA |
| *cln3(4m)-MoonTag* | TAATACGACTCACTATAAGGGACGTCCTAGTACCAAGGATCAGCCGCTCTAGTTAAAGGGGACGAACCGGGGCTCTTATCCTTAGCATCGTCTCTTGTCCACACCTCGTTGACCTGCAGACACAGCACAAACCCCTAGTAATTAATAGTAATCATTGTCATCTCCAGCTGGGCTGTTAATATCTCATACCCGAACACCTTCTAGTATTATACACATATTCGTCGCCACCATGCGTACGGGCAAGAATGAGCAGGAACTGCTGGAACTGGACAAATGGGCAAGCCTGGGGAGCGGCTCCGGAAAGAACGAGCAGGAACTGCTGGAACTGGACAAGTGGGCTTCTCTGGGCTCCGGGTCCGGGAAAAACGAACAGGAGCTGCTGGAGCTGGATAAGTGGGCCAGCCTGGGCTCCGGGTCCGGGAAGAATGAACAGGAACTGCTGGAACTGGATAAATGGGCCTCTCTGGGCAGCGGGTCCGGAAAGAATGAACAGGAGCTGCTGGAACTGGACAAGTGGGCGTCCCTGGGGAGCGGGTCCGGAAAGAATGAACAGGAACTGCTGGAACTGGATAAGTGGGCTTCCCTGGGAAGCGGCTCCGGAAAAAACGAGCAGGAGCTGCTGGAGCTGGACAAGTGGGCCTCTCTGGGCTCCGGCTCCGGGAAGAATGAACAGGAGCTGCTGGAACTGGATAAGTGGGCCTCTCTGGGGTCCGGAAGCGGTAAAAACGAGCAGGAACTGCTGGAACTGGACAAGTGGGCAAGCCTGGGGTCTGGCTCCGGCAAAAACGAACAGGAACTGCTGGAACTGGACAAATGGGCCAGCCTGGGCTCCGGGTCCGGGAAAAATGAGCAGGAACTGCTGGAACTGGATAAATGGGCGAGCCTGGGCAGCGGGTCCGGAAAAAACGAACAGGAGCTGCTGGAGCTGGACAAATGGGCGAGTCTGGGCTCCGGAAGCGGCAAGAATGAACAGGAACTGCTGGAGCTGGACAAATGGGCCAGCCTGGGCAGCGGCTCCGGAAAGAATGAGCAGGAACTGCTGGAGCTGGACAAATGGGCGTCTCTGGGCTCCGGCTCCGGGAAAAACGAGCAGGAACTGCTGGAGCTGGACAAGTGGGCTTCCCTGGGCAGCGGCTCCGGAAAAAATGAACAGGAACTGCTGGAACTGGACAAGTGGGCCAGCCTGGGGTCCGGGTCTGGGAAGAACGAGCAGGAACTGCTGGAGCTGGACAAGTGGGCTTCTCTGGGGTCTGGCTCTGGCAAGAACGAGCAGGAACTGCTGGAACTGGACAAGTGGGCGAGCCTGGGCAGCGGGTCTGGCAAGAATGAACAGGAGCTGCTGGAACTGGACAAATGGGCCAGCCTGGGGTCTGGGTCTGGAAAGAATGAGCAGGAACTGCTGGAGCTGGACAAGTGGGCCTCTCTGGGCTCTGGGTCTGGCAAGAACGAGCAGGAGCTGCTGGAGCTGGATAAGTGGGCGTCCCTGGGGTCCGGGTCCGGCAAGAATGAACAGGAGCTGCTGGAGCTGGATAAATGGGCTTCCCTGGGCTCTGGGTCTGGGAAGAATGAGCAGGAGCTGCTGGAACTGGATAAGTGGGCGAGCCTGGGCAGCGGGTCTGGCAAGAATGAGCAGGAACTGCTGGAGCTGGACAAGTGGGCTTCTCTGTAA |
| *BNI1-MoonTag* | TAATACGACTCACTATAAGACTAACATATAGAGGGCAGTACAGCACACGTCAACTATCTTTGCTCGTCGATCGTCGGGCATTTACACACCACAGTACACGCGCGCGCCTGGGGCTTGGAGATACATTTCTGCTTGCAGGCCCGAATATTGCGCTAATCGTCAAGGCACTCCAGCATTCCACGCAGTGGGACAAGGCAGCAGGTGTGGAACACGGCGGGACGCAGGATTTACCGGTATAGTGGCAGCGTCGGCGGCTGGGCACATATGCAAGGGCCACCATGAAGAAGTCCACGCACTCGAACAAACACTCCAAAACACAGCACGCGGGGCAGGTTGCCGGGGCAACACACGCCAACGGTGGCCACAGCGGCGGTGGCTTGTTCTCGAACCTCATGAAGTTGACGGGCTCGTCGGGCTCGCAGAGGATTGCGACCTCGGATATTTCGTCTCCGAAGAAGGTGACGCTGCCGTCCAATCTGGACCTGTCCGGTGCAAAACCCTTGAACAAGCTGAATACGATCAACGCGTCCAATTTACCACAGGTGTCGGGCGAGGTAACACAGGCCCGATCTGCGTCTGGCGCTGCGTTGTCTTCACCATCCAAGTATTCATATTCCAAGCGGTCTTCCCAGTGGGCGAGCTCCAAGCAAGCAAACCATCCAGAACACGACCAGCAGTATCCCCAGCATCTTAACCATCTTTTTGTGCAGCCCCATCCTCACCACTCGGCTTTGACGCGGCAGATGACCAACCAATCAAATTACTCCGCTTCTTCATATACTTCTTTATCTCTGCTACATAAAGCCACGGATATAGATGGTACCTTAACCTTGGAGAAGCCCGAGGATCCCCAAGAGATTGAAGAACTATACCAGGAGTTGCTACAGAAAAGAAACATACCACAATCCGTTTCTGTCCACGGTCATCGCGAGCTGATGAGTTATGGCATTGATAAAAAGTGGCTTATGGTTAAACAGGATCTCCAAACGGAATACAAGAAGATGAAAAATAGCATGCCACCAAGCTCGACCGTAGAGTCGATAGCGAGTTCTGCTCTAGCAAAATCCCCTGGCGGAAGCATATCGTCCGGGAGCATCACAAGATCCATATCTCAAAACAGCGCGAAGCCCATGGCTGTAGCCTACGAGCCTGCACATCTCTCGCCCGATCACTACGTGCGGAAGATTATATCGGATAGAGTGACTGCACAAGAACTAAACGATTTGTGGATTTCGTTGAGGACCGAAAGCATAGACTGGGTGATTGGTTTTATCGACGCTCAGGGACAGGTTGCAATTGCCAATCGCCTTTTAAAATTTGTGCAACGTGAGTCTCTAGATATTCTACATGATTATGATGCATTAGAGAAGGAAAACGCCTACTATAAGTGTTTGAGAGTCCTTACGAACTTGCGGGAAGGGATGCAGGAAGCGCTGAAGTCCAAACTAGTGATCAGCTCTGTGGTGGAGGGTCTATTATCCAGTAGGTTGTCCACCAGGAGGGTAGCAACGGAGACTCTATTGTACATGCTCTCTGACGACGAGCTCAAAACTGCAGATACGGGTCAAGATGCGATTTTTCCTTTGTTGGTTGCATTAGACCAAGAATCAAAGTTTGCTGCAAACATTCATCTTCGTGGAAGACTTCATGAAACTAATAAACGTAAATCCTTACCTACTGGGAGAGCAGATTTCCATGTGGTCGCCAAACGACTAGAACAATGGCTATATGTCATTGAATATACCTTGGACGGTCGCGGGAGAATGGGTTCTCTTGTTGGTGCATCTGGTACGGGCAAGAATGAGCAGGAACTGCTGGAACTGGACAAATGGGCAAGCCTGGGGAGCGGCTCCGGAAAGAACGAGCAGGAACTGCTGGAACTGGACAAGTGGGCTTCTCTGGGCTCCGGGTCCGGGAAAAACGAACAGGAGCTGCTGGAGCTGGATAAGTGGGCCAGCCTGGGCTCCGGGTCCGGGAAGAATGAACAGGAACTGCTGGAACTGGATAAATGGGCCTCTCTGGGCAGCGGGTCCGGAAAGAATGAACAGGAGCTGCTGGAACTGGACAAGTGGGCGTCCCTGGGGAGCGGGTCCGGAAAGAATGAACAGGAACTGCTGGAACTGGATAAGTGGGCTTCCCTGGGAAGCGGCTCCGGAAAAAACGAGCAGGAGCTGCTGGAGCTGGACAAGTGGGCCTCTCTGGGCTCCGGCTCCGGGAAGAATGAACAGGAGCTGCTGGAACTGGATAAGTGGGCCTCTCTGGGGTCCGGAAGCGGTAAAAACGAGCAGGAACTGCTGGAACTGGACAAGTGGGCAAGCCTGGGGTCTGGCTCCGGCAAAAACGAACAGGAACTGCTGGAACTGGACAAATGGGCCAGCCTGGGCTCCGGGTCCGGGAAAAATGAGCAGGAACTGCTGGAACTGGATAAATGGGCGAGCCTGGGCAGCGGGTCCGGAAAAAACGAACAGGAGCTGCTGGAGCTGGACAAATGGGCGAGTCTGGGCTCCGGAAGCGGCAAGAATGAACAGGAACTGCTGGAGCTGGACAAATGGGCCAGCCTGGGCAGCGGCTCCGGAAAGAATGAGCAGGAACTGCTGGAGCTGGACAAATGGGCGTCTCTGGGCTCCGGCTCCGGGAAAAACGAGCAGGAACTGCTGGAGCTGGACAAGTGGGCTTCCCTGGGCAGCGGCTCCGGAAAAAATGAACAGGAACTGCTGGAACTGGACAAGTGGGCCAGCCTGGGGTCCGGGTCTGGGAAGAACGAGCAGGAACTGCTGGAGCTGGACAAGTGGGCTTCTCTGGGGTCTGGCTCTGGCAAGAACGAGCAGGAACTGCTGGAACTGGACAAGTGGGCGAGCCTGGGCAGCGGGTCTGGCAAGAATGAACAGGAGCTGCTGGAACTGGACAAATGGGCCAGCCTGGGGTCTGGGTCTGGAAAGAATGAGCAGGAACTGCTGGAGCTGGACAAGTGGGCCTCTCTGGGCTCTGGGTCTGGCAAGAACGAGCAGGAGCTGCTGGAGCTGGATAAGTGGGCGTCCCTGGGGTCCGGGTCCGGCAAGAATGAACAGGAGCTGCTGGAGCTGGATAAATGGGCTTCCCTGGGCTCTGGGTCTGGGAAGAATGAGCAGGAGCTGCTGGAACTGGATAAGTGGGCGAGCCTGGGCAGCGGGTCTGGCAAGAATGAGCAGGAACTGCTGGAGCTGGACAAGTGGGCTTCTCTGTAA |
| *bni1(3m)-MoonTag* | TAATACGACTCACTATAAGACTAACATATAGAGGGCAGTACAGCACACGTCAACTATCTTTGCTCGTCGATCGTCGGGCATTTACACACCACAGTACACGCGCGCGCCTGGGGCTTGGAGATACATTTCTGCTTGCAGGCCCGAATATTGCGCTAATCGTCAAGGCACTCCAGCATTCCACGCAGTGGGACAAGGCAGCAGGTGTGGAACACGGCGGGACGCAGGATTTACCGGTATAGTGGCAGCGTCGGCGGCTGGGCACATATGCAAGGGCCACCATGAAGAAGTCCACGCACTCGAACAAACACTCCAAAACACAGCACGCGGGGCAGGTTGCCGGGGCAACACACGCCAACGGTGGCCACAGCGGCGGTGGCTTGTTCTCGAACCTCATGAAGTTGACGGGCTCGTCGGGCTCGCAGAGGATTGCGACCTCGGATATTTCGTCTCCGAAGAAGGTGACGCTGCCGTCCAATCTGGACCTGTCCGGTGCAAAACCCTTGAACAAGCTGAATACGATCAACGCGTCCAATTTACCACAGGTGTCGGGCGAGGTAACACAGGCCCGATCTGCGTCTGGCGCTGCGTTGTCTTCACCATCCAAGTATTCATATTCCAAGCGGTCTTCCCAGTGGGCGAGCTCCAAGCAAGCAAACCATCCAGAACACGACCAGCAGTATCCCCAGCATCTTAACCATCTTTTTGTGCAGCCCCATCCTCACCACTCGGCTTTGACGCGGCAGATGACCAACCAATCAAATTACTCCGCTTCTTCATATACTTCTTTATCTCTGCTACATAAAGCCACGGATATAGATGGTACCTTAACCTTGGAGAAGCCCGAGGATCCCCAAGAGATTGAAGAACTATACCAGGAGTTGCTACAGAAAAGAAACATACCACAATCCGTTTCTGTCCACGGTCATCGCGAGCTGATGAGTTATGGCATTGATAAAAAGTGGCTTATGGTTAAACAGGATCTCCAAACGGAATACAAGAAGATGAAAAATAGCATGCCACCAAGCTCGACCGTAGAGTCGATAGCGAGTTCTGCTCTAGCAAAATCCCCTGGCGGAAGCATATCGTCCGGGAGCATCACAAGATCCATATCTCAAAACAGCGCGAAGCCCATGGCTGTAGCCTACGAGCCTGCACATCTCTCGCCCGATCACTACGTGCGGAAGATTATATCGGATAGAGTGACTGCACAAGAACTAAACGATTTGTGGATTTCGTTGAGGACCGAAAGCATAGACTGGGTGATTGGTTTTATCGACGCTCAGGGACAGGTTGCAATTGCCAATCGCCTTTTAAAATTTGTGCAACGTGAGTCTCTAGATATTCTACATGATTATGACGCTTTAGAGAAGGAAAACGCCTACTATAAGTGTTTGAGAGTCCTTACGAACTTGCGGGAAGGGATGCAGGAAGCGCTGAAGTCCAAACTAGTGATCAGCTCTGTGGTGGAGGGTCTATTATCCAGTAGGTTGTCCACCAGGAGGGTAGCAACGGAGACTCTATTGTACATGCTCTCTGACGACGAGCTCAAAACTGCAGATACGGGTCAAGATGCGATTTTTCCTTTGTTGGTAGCCTTAGACCAAGAATCAAAGTTTGCTGCAAACATTCATCTTCGTGGAAGACTTCATGAAACTAATAAACGTAAATCCTTACCTACTGGGAGAGCAGATTTCCATGTGGTCGCCAAACGACTAGAACAATGGCTATATGTCATTGAATATACCTTGGACGGTCGCGGGAGAATGGGTTCTCTTGTTGGGGCTTCTGATTGTACGGGCAAGAATGAGCAGGAACTGCTGGAACTGGACAAATGGGCAAGCCTGGGGAGCGGCTCCGGAAAGAACGAGCAGGAACTGCTGGAACTGGACAAGTGGGCTTCTCTGGGCTCCGGGTCCGGGAAAAACGAACAGGAGCTGCTGGAGCTGGATAAGTGGGCCAGCCTGGGCTCCGGGTCCGGGAAGAATGAACAGGAACTGCTGGAACTGGATAAATGGGCCTCTCTGGGCAGCGGGTCCGGAAAGAATGAACAGGAGCTGCTGGAACTGGACAAGTGGGCGTCCCTGGGGAGCGGGTCCGGAAAGAATGAACAGGAACTGCTGGAACTGGATAAGTGGGCTTCCCTGGGAAGCGGCTCCGGAAAAAACGAGCAGGAGCTGCTGGAGCTGGACAAGTGGGCCTCTCTGGGCTCCGGCTCCGGGAAGAATGAACAGGAGCTGCTGGAACTGGATAAGTGGGCCTCTCTGGGGTCCGGAAGCGGTAAAAACGAGCAGGAACTGCTGGAACTGGACAAGTGGGCAAGCCTGGGGTCTGGCTCCGGCAAAAACGAACAGGAACTGCTGGAACTGGACAAATGGGCCAGCCTGGGCTCCGGGTCCGGGAAAAATGAGCAGGAACTGCTGGAACTGGATAAATGGGCGAGCCTGGGCAGCGGGTCCGGAAAAAACGAACAGGAGCTGCTGGAGCTGGACAAATGGGCGAGTCTGGGCTCCGGAAGCGGCAAGAATGAACAGGAACTGCTGGAGCTGGACAAATGGGCCAGCCTGGGCAGCGGCTCCGGAAAGAATGAGCAGGAACTGCTGGAGCTGGACAAATGGGCGTCTCTGGGCTCCGGCTCCGGGAAAAACGAGCAGGAACTGCTGGAGCTGGACAAGTGGGCTTCCCTGGGCAGCGGCTCCGGAAAAAATGAACAGGAACTGCTGGAACTGGACAAGTGGGCCAGCCTGGGGTCCGGGTCTGGGAAGAACGAGCAGGAACTGCTGGAGCTGGACAAGTGGGCTTCTCTGGGGTCTGGCTCTGGCAAGAACGAGCAGGAACTGCTGGAACTGGACAAGTGGGCGAGCCTGGGCAGCGGGTCTGGCAAGAATGAACAGGAGCTGCTGGAACTGGACAAATGGGCCAGCCTGGGGTCTGGGTCTGGAAAGAATGAGCAGGAACTGCTGGAGCTGGACAAGTGGGCCTCTCTGGGCTCTGGGTCTGGCAAGAACGAGCAGGAGCTGCTGGAGCTGGATAAGTGGGCGTCCCTGGGGTCCGGGTCCGGCAAGAATGAACAGGAGCTGCTGGAGCTGGATAAATGGGCTTCCCTGGGCTCTGGGTCTGGGAAGAATGAGCAGGAGCTGCTGGAACTGGATAAGTGGGCGAGCCTGGGCAGCGGGTCTGGCAAGAATGAGCAGGAACTGCTGGAGCTGGACAAGTGGGCTTCTCTGTAA |

### *In vitro* transcription

*In vitro* transcription (IVT) of translation-competent *Nluc* or *MoonTag* mRNAs were carried out according to manufacturer’s specifications (NEB, E2080S or E2040S). cDNA template was amplified using PCR for all Nluc constructs. For the production of *MoonTag-containing* RNA, plasmids were linearized with HindIII digestion to generate cDNA template. PCR products (or digestion reactions) were purified with a PCR purification kit (Qiagen, 28104) before IVT. IVT was carried out using HiScribe T7 with CleanCap Reagent AG (NEB, E2080S) to produce Cap-1 *mRNA*. These reactions did not include Cy5-UTP, to reduce the impact of bulky adducts on translation. Following the first purification and precipitation with 2.5 M LiCl, RNA poly(A) tails were added to the Cap-1 mRNAs according to manufacturer specifications (NEB, M0276L). Reactions were then purified a second time with 2.5 M LiCl precipitation. RNA concentration was determined by 260 nm absorbance and verified for purity and size using a denaturing agarose gel and Millenium RNA ladder (ThermoFisher Scientific, AM7151).

For Cy5 labeled RNAs, IVT was carried out according to manufacturer’s instructions (NEB, E2040S), with the following modifications. Reactions included 1 µg of DNA template and were supplemented with approximately 0.4 µl of Cy3-labelled UTP (Sigma, PA53026). RNA concentration was determined by 260 nm absorbance and verified for purity and size as described above.

To determine RNA labelling efficiency, following absorbance measurements at 260 nm, the same RNA was measured for Cy5 absorbance at 650, with an extinction Coefficient of 250000 (l/mole-cm), and corrected for absorbance at 260 and 280 using correction factors 0.00, and 0.05 respectively. The labelling efficiency was then derived using the formula:

$$\frac{RNA concentration (\mu M)}{Cy5 concentration (\mu M)}$$

Labelling efficiency was used to normalize grey values acquired in the Cy5 channel.

| Primer | Sequence |
| --- | --- |
| BNI1-Nluc_fwd | GAAAATTGTCCCAATTAGTAGCATCACGCT |
| bni1(3m)-Nluc_fwd | GAAAATTGTCCCAATTAGTAGCATCACGCT |
| CLN3-Nluc_fwd | ttcgtgCACGCTGTGtaatacgactcactataagggtctgcataccaaggat |
| cln3(4m)-Nluc_fwd | ttcgtgCACGCTGTGtaatacgactcactataagggacgtcctagtaccaag |
| Nluc_rev | ttatttgtacaatgatcccccacccgc |

### Preparation of imaging chambers

For in vitro condensate studies, #1.5 glass coverslips (22x22; VWR 48393-230) were treated with oxygen plasma for 10 min to clean the glass. After plasma cleaning, imaging wells were created using the tops of 0.2 mL tubes glued to the coverslip surface using UV-cured glue (Norland Optical Adhesive, NOA 60). Imaging wells were than loaded with 25 µl of 1.5 mg/mL PLL-g-PEG dissolved in ddH_2_O (SuSoS SZ40-89) and left to incubate for 1 hr at RT in a humidity-controlled environment. After incubation, imaging wells were rinsed 4x with 50 µl ddH_2_O before being filled with 50 µl translation buffer (30mM Hepes (pH 7.4; KOH), 120mM Potassium Glutamate and 2mM Magnesium Glutamate) until use. Reaction chambers emptied of buffer before use and were used on the day of passivation.

### Labeling ribosomes for *in vitro* reactions

E. coli ribosomes (NEB; P0763S) were labeled with Atto 565 NHS ester (Supelco; 72464) by combining the whole ribosome aliquot with 3x molar excess Atto dye. The reaction was gently mixed by pipetting, wrapped in aluminum foil and left at RT for 30 minutes away from light. After labeling, the volume of the reaction was adjusted to 0.5 mL using reaction buffer (50 mM Hepes at pH 7.4, 150 mM KCl) and loaded into a Amicon Ultra 0.5 mL centrifugal filter (MWCO 30 kDa; Millipore UFC5030). Ribosomes were purified from the unreacted dye according to manufacturer’s specifications (Millipore; UFC5030) by repeating five cycles of spin concentration followed by resuspension in clean reaction buffer.

In vitro reactions containing ribosomes were supplemented with 400 nM labeled ribosomes immediately after ribosomes were thawed and labeled.

### Fluorescence microscopy of condensates

3D fluorescence imaging of in vitro condensate reactions was performed on a Nikon Ti2-E inverted microscope equipped with a Yokogawa CSU-W1 spinning disk confocal and an ORCA-Fusion BT Digital CMOS camera. Images were collected using a 100x oil objective (CFI Plan Lambda, NA = 1.45, WD = 0.13 mm) with Nyquist sampling.

### Preparation of Ashbya lysates

*Ashbya* lysates were prepared using a protocol adapted from a previously established yeast cell-free extract protocol ^2^. Cells were allowed to grow overnight for 16 hrs prior to their collection via vacuum filtration. Mycelial mats were rinsed with RT PBS (1x; 500 mL of Ashbya culture to 150 mL of PBS) to remove residual media. After washing the mycelial mat, mycelia were gently scraped into a pre-cooled mortar and pestle containing liquid N_2_. After flash freezing, the cells were ground into a fine powder, ensuring that the mortar and pestle contained liquid N_2_ throughout the grinding process, reducing temperature fluctuations during cell lysis. After mycelia were ground, the remaining liquid N_2_ was volatilized and the dry mass of the sample was measured, and for every 1 g of dry mass, 300 µl of ashbya-lysis buffer (50 mM Hepes-KOH, 140 mM KCl, 1 mM MgCl_2_, 1 mM EGTA, 0.1 % Tween-20, 5% glycerol), 4 uL of protease inhibitor cocktail (Millipore Sigma, 539136-1ML), and 40 U of Superase-In RNAse inhibitor (ThermoFischer, AM2696) were added to the dry-cell mass. RNAse inhibitor was excluded from lysates treated with RNAse A (see “Proteomics”). The ground mycelial was allowed to thaw on ice with gentle mixing every 15 min. Crude lysates were transferred to centrifuge-safe Eppendorf tubes (Beckman Coulter, 357448) and centrifuged at 70,000 RPM in a TLA 100.3 rotor (112,000 RCF) for 20 min. Supernatant was collected with care taken not to disturb the cell pellet. The spin step was repeated once more, and the cell-free extract collected was used the day of its preparation.

### Preparation of SUVs

SUVs were prepared as previously described in ^3^, with the following modifications. SUVS were prepared using 96 mol% DOPC and 4 mol% DGS NTA-Ni (Avanti Polar Lipids: 850856C and 790408C). After mixing lipids in chloroform solvent at the specified molar ratios, the mixture was dried by a stream of nitrogen gas to generate a lipid film. The remaining chloroform was evaporated by placing the lipid film under a vacuum for at least 2 h. Lipids were rehydrated in 50 mM Hepes (pH 7.4) and 150mM KCl to a final concentration of 0.5-1mM. Hydrated lipids were subject to bath sonication (Symphony) until clarification to yield small unilamellar vesicles (SUVs). SUVs were aliquoted and stored at -80 until use. SUVs in this study were used within one month of their preparation.

### Proteomics

First, SUVs were adsorbed onto 3 um (ø) silica microspheres as previously described ^4^. Briefly, 50 nM lipids were added to reaction buffer (50 mM Hepes (pH 7.4) and 150mM KCl) for every 440 µm^2^ of silica microsphere surface area in the reaction. Reactions were left for 1 h on a roller drum at RT. Unbound lipid was washed away by pelleting lipid-coated beads at 800 RCF and removing unbound lipids. Beads were then washed using reaction buffer. Washes were performed four times for a total of 8x reaction volumes. Beads were then resuspended in 100 µl of reaction buffer. Recombinantly expressed Whi3 was added to the resuspended beads to a final concentration of 4 µM and left to incubate at RT for 30 min on a roller drum. Subsequently, the beads were pelleted at 800 RFC, and unbound Whi3 was removed by resuspending and washing the beads four times in reaction buffer for a total of 8x reaction volumes. The beads were resuspended to a final volume of 60 µl.

For each treatment (+RNAse A and -RNAse A), three separate Ashbya lysates were prepared (see “Preparation of Ashbya cell-free extracts”). For each biological replicate (each separate lysate), the lysate was split into two 540 µl fractions. One fraction was supplemented with 60 µl Whi3-coated silica microspheres while the other fraction was supplemented with 60 µl lipid-coated silica microspheres without Whi3. These reactions were left to incubate for 1.5 hr on a roller drum at 30°C. After incubation in CFE, the beads were pelleted at 800 RCF and as much CFE was removed without disturbing the bead pellet. The bead pellet was than washed 3x with 500 µl of reaction buffer. The pellet was subsequently resuspended in 50 µl elution buffer (50 mM Hepes (pH 7.4), 150 mM KCl, 200 mM Imidazole) and left to incubate at RT for 10 minutes. The beads were pelleted at 800 RCF and the supernatant was carefully removed. The eluted sample containing either Whi3- or control-bound proteins was then submitted to the UNC Proteomics core for AP-MS/MS analysis. Importantly, comparisons of protein enrichment between Whi3-coated beads and lipid-coated beads (control) were limited to reactions that originated from the same extract. Additionally, in order to be considered enriched across the three extracts, the unique protein IDs had to (i) be found in at least two of the treatment samples, (ii) have a P-value of ≤ 0.05 when compared to the control sample from the same extract by a Students T-test, and (iii) a log2(FC) ≥ 1.5 (3x more abundant) by the same Students T-test when compared to the control sample. For RNAse A treated Ashbya CFEs, CFEs were supplemented with 100 µg/ml RNAse A (ThermoFischer; EN0531) and incubated at 30°C for 30 min prior to the addition of Whi3-coated microspheres.

Gene ontology (GO) analysis was carried by using the publicly available g:Profiler (<https://biit.cs.ut.ee/gprofiler/orth>), which compared the enriched proteins found in our data set to those genes contained within ATCC 10895 (*Ashbya gossypii*) by cellular compartment. After GO analysis, results were loaded into Python 3.11.3 for data wrangling and visualization. For scatter plot visualization, genes associated with the following GO terms were colored light blue for translation-association: GO:0006412, GO:0002181. Data points associated with the following GO terms were colored light orange for RNA regulation; GO:0006402, GO:0006401, GO:0004535.

### Assembly of condensates

To assemble Whi3, whi3(S637D), whi3(ΔQRR), Fus, or N-protein condensates, protein and RNA were added to translation buffer (30mM Hepes (pH 7.4; KOH), 120mM Potassium Glutamate and 2mM Magnesium Glutamate) at 2x the final concentration. Unless specified, final concentrations were 3 µM protein and 5 nM RNA. These reactions were allowed to incubate at room temperature for 10 minutes before the addition of cell-free lysate to a final concentration of 1:1 (i.e. 10 µl of RR lysate to 10 µl condensate mix), bringing the final reaction concentrations of Whi3 and RNA to 1x final concentration. For fluorescence imaging, unlabeled protein was mixed with Atto488 labeled protein such that the final labeled protein fraction was equal to 5% for all reactions.

Condensates formed in the presence of ribosomes were allowed to mature for 10 min prior to the addition of 400 nM ribosomes-atto565.

### Luciferase assay

For luciferase assays, 5 nM *CLN3-Nluc*, *cln3(4m)-Nluc*, *BNI1-Nluc*, or *bni1(3m)-Nluc,* and increasing concentrations (0, 50, 100, 1000, and 3000 nM) of Whi3, whi3(S637D), or whi3(ΔQRR) were pre-mixed in translation buffer (30mM Hepes (pH 7.4; KOH), 120mM Potassium Glutamate and 2mM Magnesium Glutamate) to ½ the total reaction volume and left to incubate for 10 min at RT prior supplementing the reaction 1:1 with RR extract (Promega, L4960). The reactions were incubated at 30°C for 1 hr before the addition of substrate, Vivazine (Promega, N2580). Vivazine was diluted 1:50 in translation buffer, and 15 µl of substrate was added for every 20 µl of luciferase reaction. The luminescence for each reaction was read out on a Promega GloMax® Explorer with a 2 s integration time.

For luminescence data wrangling, visualization and statistical analysis, raw luminescence values were read into Python 3.11.3. To minimize the impact of lysate-to-lysate variation on luminescence values, the analysis was carried out as follows: for each condition (0, 50, 100, 1000, or 3000 nM Whi3), data were collected from at least three technical replicates per an extract, for a total of three extracts or biological replicates. Luminescence values were than normalized and expressed as log_2_(FC) to 0 nM Whi3 conditions that occurred within the same biological lysate.

For luciferase reactions that underwent centrifugation, reactions were prepared as described above with the following modifications: After the addition of mRNA and Whi3 to translation buffer, the reactions were immediately centrifuged at 500 or 10000 RCF. The luminescence associated with these reactions were normalized and expressed as log_2_(FC) over reactions containing 0 nM Whi3 but that were spun at identical speeds for the same duration. As before, data were collected from at least three technical replicates per lysate, for a total of three lysates.

### RNA turnover and decay

To measure RNA turnover in our luciferase reactions, Cy5-labeled RNA and either buffer or 3 µM Whi3 were mixed as described in “Assembly of Condensates” and supplemented with RR lysate. Reactions were loaded onto a 96 well plate and measured for Cy5 signal upon the addition of RR lysate (0 hr) and after one hour at 30C (1 hr). Cy5 signal was measured on a Promega GloMax® Explorer with an excitation wavelength of 627 nm and emission wavelength of 660-720, using the “Highly Sensitive” optimization setting.

### Measuring condensate area

To measure condensate area associated with different concentrations of Whi3 and mRNA, condensate-forming reactions were carried out as described above (see “Assembly of condensates”) and loaded into imaging chambers (see “Preparation of imaging chambers”). After approximately 20 min, reactions were subject to 3D fluorescence imaging (see “Fluorescence microscopy”).

After acquisition, condensate number, area, and area fraction were measured using a custom macro in Fiji. The macro performed the following operations: Each individual image-stack was first cropped to the center 1200x1200 pixels of the camera chip (total camera chip = 2304x2304 pixels) to minimize uneven illumination across the field of view. Image stacks were then projected as a sum intensity. Condensates were segmented out of the image by (i) duplicating the image, (ii) thresholding via automated Otsu’s method, and (iii) using particle detection on the mask. After particle detection, the results were read into Python 3.11.3 for wrangling and data visualization. Condensate areas are reported from three fields of view (FOV) for each of the three separate RR lysates tested for each condition.

### Measuring partitioning coefficients and volume fractions of Whi3 condensates

To estimate dense and dilute phase concentrations in our minimal system, condensate-forming reactions were carried out as described above (see “Assembly of condensates”) and loaded into imaging chambers (see “Preparation of imaging chambers”). After approximately 20 min, reactions were subject to 3D fluorescence imaging (see “Fluorescence microscopy”).

After acquisition, we first estimated volume fraction using a custom macro in Fiji. The macro performed the following operations: Each individual image-stack was first cropped to the center 1200x1200 pixels of the camera chip (total camera chip = 2304x2304 pixels) to minimize uneven illumination across the field of view. Condensates were segmented from the image stack by (i) duplicating the stack, (ii) thresholding via automated Otsu’s method, and (iii) using particle detection on the mask. After particle detection, the results were read into Python 3.11.3 for wrangling and data visualization. To estimate volume fraction per image slice, the total area of the image and the area occupied by the condensates were multiplied by the voxel height. Volume fraction was estimated by dividing the total volume occupied by condensates by the total volume of the image stack. Volume fractions were measured from three fields of view (FOV) for each of the three separate RR lysates tested for each condition.

To estimate partitioning coefficient (Kp), individual image-stacks were first cropped to the center 1200x1200 pixels of the camera chip (total camera chip = 2304x2304 pixels) to minimize uneven illumination across the field of view. Image stacks were then projected as a sum intensity. Condensates were segmented out of the image by (i) duplicating the image, (ii) thresholding via automated Otsu’s method, and (iii) using particle detection on the mask. After particle detection, Whi3 and RNA intensities per condensate were read into Python 3.11.3 for wrangling and data visualization. Partitioning coefficient was calculated by dividing the mean intensity of the condensate by the background intensity within the same image.

Dense and dilute phase concentrations of Whi3 and RNA could then be calculated using the following equations ^5^:

$$Equation 1: C_{total}= C_{dense}V_{f}+C_{dilute}\left( 1-V_{f} \right)$$

$${Equation 2: C}_{dense}= K_{p}C_{dilute}$$

C_total_= initial concentration

V_f_= volume fraction of the dense phase

K_p_= partition coefficient

### MoonTag in extract

The MoonTag approach was modified from existing protocols ^6^ to work in RR lysates. Condensates were formed as previously described (see “Assembly of condensates”) and allowed to incubate in RR lysates containing 500 nM 2H10-mCherry for 1 hr at 30C prior to being loaded into imaging chambers (see “Preparation of imaging chambers”). After approximately 20 min, reactions were subject to 3D fluorescence imaging (see “Fluorescence microscopy”). For cycloheximide experiments, Ashby lysates were supplemented with 100 µg/ml cycloheximide dissolved in 50 mM Hepes (p.H. 7.4; KOH) and 150 mM KCl.

Image analysis of Whi3, whi3(S637D), Fus, and N-protein condensates and the associated MoonTag signal associated with each condensate was carried out using a custom macro in Fiji. The macro performed the following operations: Each image-stack was first cropped to the center 1200x1200 pixels of the camera chip (total camera chip = 2304x2304 pixels) to minimize uneven illumination across the field of view. Image stacks are then projected as a sum intensity, and the median value of the pixels corresponding to the MoonTag channel (closely approximating the background intensity of soluble MoonTag signal) was subtracted from the MoonTag channel. A duplicate image corresponding to the RNA-binding protein channel is generated for automatic image thresholding via Otsu’s method, a one pixel-dilation on the threshold mask, and subsequently particle detection on the mask to generate a list of ROIs corresponding to the condensates. The ROIs corresponding to the RBP condensates were then overlaid on the fluorescence channels corresponding to condensate and MoonTag channels, and the average fluorescence intensity values for each ROI were collected.

Raw fluorescence values extracted from Fiji were then loaded into Python 3.11.3 for wrangling, statistical analysis, and data visualization. For each condition or RBP, data were collected from three RR lysates, or biological replicates. For each lysate, at least three image stacks were collected for analysis.

### Radial intensity profiles

Plot profiles used to represent the distribution of fluorescence intensities corresponding to MoonTag translation, ribosomes, or RNA with respect to Whi3, whi3(S637D), Fus, or N-protein condensates were generated using a custom Fiji and Python script. To reduce variation in condensate-midplane for condensates of different sizes contained within the same FOV, images were first cropped to generate single-condensate image stacks (or regions of similar sized condensates). Image stacks were then manually analyzed for the 2D-cross section which most closely approximated the midplane of the condensates. After midplane selection, the custom Fiji macro performed the following operations: For each image, a list of ROIs corresponding to the condensates was generated by (i) duplicating the RBP channel, (ii) thresholding via automated Otsu’s method to generate a mask, and (iii) using particle detection on the mask to generate ROIs. For each ROI in the list of ROIs, the centroid and largest dimension of the bounding box corresponding to that ROI were used to identify the center of the condensate and the radius of the condensate, respectively. Using the plugin, “Radial Profile”, populated with the x,y coordinate positions corresponding the condensate centroid, with a radius of analysis equal to 10x the radius of the condensate, a plot profile was generated for each ROI, for each channel.

After collection in Fiji, radial plot profiles for all condensates measured were loaded into Python 3.11.3 for data wrangling and analysis. To allow for the superimposition of radial plot profiles from condensates of different size and fluorescence intensity, we used a custom Python script that performed the following operations: for each condensate, the radius of analysis (similar to ROI length, see “Radial Profile” description) was normalized by the maximum distance of that radial profile, such that the maximum radial position is now 1, with the condensate radius approximately equal to 0.1. Next the fluorescence intensity value associated with each radial position was normalized to the fluorescence intensity associated with the center of that condensate, or at x=0. After these normalizations, SciPy’s sub-package, scipy.interpolate, was used to impute fluorescence intensity values at defined x-positions along the radial plot profile. After interpolation, the average fluorescence intensity value at each x-position as well as the standard error of the mean associated with that fluorescence value, across all condensates, was calculated. For visualization of radial plot profiles, the average intensity value of all condensates included in the analysis is plotted at that x-position with the error corresponding to ±SEM.

For analysis of interface intensity per pixel, or for calculations of translation peak position relative to the condensate center, radial plot profiles were loaded into Python 3.11.3 as before, however individual radial profiles were not normalized. Instead, raw intensity values and pixel distances were used to calculate condensate radius, translation peak location, and intensity per pixel at the condensate interface. Condensates whose interface wasn’t resolvable from the condensate center (condensate radius < 300 nm) were indicated graphically.

### Live cell imaging

*A. gossypii* ade2:H4-GFP:ADE2 (AG368)/Whi3-tomato:NAT heterokaryons were grown by inoculating dirty spores in 25 mL AFM for 16 h in a shaking incubator at 30 ⁰C under selection at a final concentration of 75 µg/mL nourseothricin sulphate. After 16 h of growth cells were spun down at 400 rpm for 5 min. Media was removed from the pellet and they were washed with twice with 10 mL 2xLFM. Cells were then resuspended with 3 mL 2xLFM after a second wash and 20 μL of the suspension was placed on a LFM gel pad (2% agarose in 2xLFM) which was covered with a coverslip and sealed with VALAP. The cells were allowed to recover at room temperature for 1 hour before the start of imaging.

Imaging was performed on Zeiss Lattice Lightsheet 7 using a 40x/NA 1.0 water immersion objective. Hyphae were imaged under 488 nm and 561 nm excitation for 45 min at 90 s intervals between each frame. All images were taken within 1-3 hours of being placed on the microscope over multiple days to generate 21 movies in total.

For cell cycle correlations, AG270.1 (Tub4-mCherry:NAT) was transformed with plasmid AGB1396 encoding Whi3-mNeon:KAN, to make AG1028. 3 different transformants were used to make spores and were grown for 15 h at 30 ºC under double G418 and Nat selection. 1 mL culture was gently centrifuged at 1400 rpm to collect cells which were washed once in 50 mM Tris, 150 mM NaCl and directly imaged. Cells were imaged within 30 minutes of mounting. Z-stacks were acquired on a Nikon spinning-disk confocal at 300 µm intervals with sequential imaging. 2x binning was used for mCherry imaging.

### Analysis of live cell spatial Whi3 association

For growth rate associations, max intensity projections were made of each image using Zen software. Additional analysis was performed in Fiji. Individual hyphae were cropped manually. For each movie, the hypha was marked as having a Whi3:tdTomato punctum at the tip if a punctum was present within 4 μm of the tip and remained present in two or more frames. If the punctum position relative to the hyphal tip changed within the length of the movie, the time was noted and the growth rate was from the corresponding set of frames was measured as a separate event. The growth rate was calculated by measuring the difference in the length of the hypha at either the end of the movie or changing point relative to the length of the hypha at the initial frame then dividing this difference by the corresponding time interval between the frames.

For cell cycle correlations, hyphae were cropped manually from images and both MIPs and Z-stacks were used to manually determine cell cycle stage using spindle pole appearance. Simultaneously, Whi3 puncta that made contacts with the “nuclear holes” in diffuse Whi3 signal were counted and tabulated.

### FITC-ConA pulse-chase for growth rate measurement

FITC-ConA staining was performed as previously described ^7^. *Ashbya* WT, endogenously tagged Whi3 delta QRR (AG909.2) or endogenously tagged Whi3 S637D (AG854) were grown from dirty spores in 10 mL AFM for 14 hours at 30ºC. Cell pellets were collected by centrifuging cultures for 5 minutes at 300 rpm in 15 mL tubes. Cells were then resuspended and washed twice in 10 mL wash buffer containing 50 mM Tris pH 7.5,150 mM NaCl. 50 μL of 1 mg/mL FITC-ConA (Sigma C7642) stock was added to cell pellet resuspended in 2 mL of the wash buffer and shaken at room temperature for 10 minutes. Cells were washed 2X with 10 mL AFM, resuspend cells in 10 mL AFM and appropriate selection. They were allowed to grow in a baffled flask at 300 rpm at 30°C for 1 h before fixation with 1 mL 37% formaldehyde for 1 h, shaking. Cells were washed with 1X PBS, resuspended in 500 μL 1X PBS and stained with Hoechst before mounting in 10 μL Vectasheild mounting medium (Vector Laboratories H-1000-10). FITC-ConA signal was imaged using a spinning-disk confocal and growth rates were measured by measuring the length of the tip region unlabeled with FITC in Fiji.

### Tri-probe PL-RCA for ribosome association in *Ashbya*

Tri-probes were designed as described in Zeng et al. Briefly, a fasta formatted “target” .seq file containing *CLN3* CDS, *BNI1* CDS, ACT1 CDS and a “nontarget” .seq file containing the entire *Ashbya gossypii* genome, with *CLN3*, *BNI1* and ACT1 excluded were created. These files were run on Picky2.2 on a Windows machine, set to generate 40-46 nt oligos, 330 mM salt (2X SSC), target Tm-s were set 10 ºC higher than non-targets. Each of the 4-5 probes generated were split manually into two 19-21 nt probes having similar Tms. The 5’ half was treated as the padlock probe and the 3’ half as the primer. Appropriate overhangs were concatenated.

To generate the Ag18s probes a target file containing the Ag 18S ribosomal RNA copy 39 rDNA sequence was used, and 5 probes were generated of length 25-26 nt, which partially overlapped with the paralogous Hs18s regions used in the original study. A polyA spacer and splint region were concatenated, and the 3’end designed to include three phosphorothioate modified bases and an inverted dT. All probes were ordered HPLC purified from IDT.

The PL-RCA experiment was optimized for Ashbya based on the previously reported smFISH protocol. For each experiment a 20 mL AFM culture was set up in a 250 mL baffled flask (rinsed once with sterile water, and once with AFM) with ~100 µL dirty spores and any required selection, shaking at 110 rpm at 30ºC. Cells were grown for exactly 14 hours. For translation inhibition experiments, these cultures were split into 2 flasks of 10 mL each, and G418 (1mg/mL final) or rapamycin (200nM final from 1mg/mL stock dissolved in 90% ethanol, 10% tween-20) and allowed to grow for an extra 90 minutes before fixation by shaking at 30ºC at 110 rpm. Cells were fixed by adding 37% formaldehyde (used within 3 months of opening a bottle to minimize the effects of oxidation) to a final concentration of 3.7% in the culture and shaking at 110 rpm at 30ºC for 1 h. Formaldehyde was quenched by adding 10% culture volume of 2M glycine. Flasks were incubated at 25 ºC, shaking at 100 rpm for 5 minutes to quench the fixative.

Cells were then pelleted by centrifuging at 300 rpm in 15 mL falcons for 5 minutes. The pellets were washed twice with 5 mL Buffer B (300 mM sorbitol in phosphate buffer, DEPC treated). The pellet was then moved to 1.5 mL microfuge tube, washed once more with 1 mL Buffer B at 2000 rpm, and this speed was used for all subsequent wash steps. Cells were then resuspended in lysis buffer (10 µL 200 mM VRC and 100 µL 100 mg/mL Zymolyase to cell pellet in 1 mL Buffer B). Cells were digested at 37ºC, gently shaking for 45 minutes. Optimal Zymolyase digestion time was determined empirically by performing a digestion time course for maximizing smFISH spot brightness as phase darkness did not correlate with good RCA signal. Digested cells were pelleted and washed once with Buffer B before adding 1 mL 70% Ethanol, made up in DEPC water. Cells were incubated in ethanol at 4ºC for at least 28 hours, and upto 72 hours before proceeding. Lower ethanol permeabilization times were found to yield poor spot quality for RCA.

Cells were then washed twice with Wash Buffer, containing 2X SSC, 10% Formamide and incubated at room temperature for 10 minutes in Wash buffer while the hybridization mix was prepared. For 80 µL of cell pellet, 100 µL of hybridization mixture containing hybridization buffer (2X SSC, 10% formamide, 0.1% Tween-20, 100 mg/mL dextran sulphate, 1 mg/mL yeast tRNA, 2mM VRC, 200 µg/mL BSA), pooled padlock and primer probes at 1 nM per probe (1:100 dilution of 100 nM working stock), 100 nM splint probe (1:100 dilution of 10 µM working stock), the appropriate smFISH probe set (Cln3-TAMRA, Bni1-TAMRA or Bni1-Quasar670 at 2.5 nM final) was prepared. Hybridization mix was vortexed for 30 s to mix. Finally, 1:500 dilution of anti-Tubulin antibody (Biorad MCA78G) was added to Cln3 reactions, tapped to mix. This mixture was added to the cell pellet and sharply flicked 10 times to completely mix the pellet. Hybridization was performed at 37ºC for 15 h, shaking gently.

Cells were then washed with 300 μL PBSTR (0.1 U/μl RNaseOUT, 0.1% Tween-20 in 1X PBS) twice, with short 5 min incubations at 37ºC, followed by 300 μL 20 min at 37 ºC in high-salt washing buffer (4X SSC dissolved in PBSTR) once. Finally, the samples were rinsed once with 300 μL PBSTR at room temperature before proceeding to ligation.

200 μL ligation mixture (0.25 U/μL T4 DNA ligase, 0.5 mg/ml BSA, and 0.4 U/μl of RNAseOUT inhibitor in 1X T4 DNA ligase buffer) was added to the cell pellet, tapped to mix and incubated at room temperature for two hours with gentle shaking. Cells were washed twice with 300 μL PBSTR.

200 μL rolling circle amplification mixture (0.5 U/μl Phi29 DNA polymerase, 250 μM dNTP, 0.5 mg/ml BSA and 0.4U/μl of RNaseOUT in 1X Phi29 buffer) was added to cell pellets, tapped to mix, and incubated at 4 ºC for 30 min before incubating at 30 ºC for two hours with gentle shaking. Cells were washed twice with PBST (0.1% Tween-20 in PBS).

Cells were then incubated with readout probe at 100 nM final and AlexaFluor-488 labeled anti-rat secondary antibody (Abcam ab150109) 1:1000 in 200 µL of 2X SSC, including Hoechst at 1:1000 dilution of 1 mg/mL stock. Cells were washed twice with 1X PBST.

15 µL of cells were mount in 20 µL of Vectashield mounting medium by pipetting thrice slowly to mix and immediately placing under a coverslip. Cells were flattened by placing under a 500 mL glass bottle for 20 minutes. The edges were sealed using Xtreme Wear transparent nail polish and allowed to air dry in the dark for 5-10 minutes. Faster drying times were correlated with brighter signal. Slides were stored at -20ºC, until they were imaged, typically within 24 hours of mounting.

For Whi3 immunofluorescence, cells expressing Whi3-mNeon (AG1028.2-4, generated by transforming plasmid AGB1396, pRS416-Whi3-mNeon, into AG270) were used as described previously for PL-RCA. Additionally, the primary anti-mNeon antibody 32F6 (ChromoTek, Proteintech) was included at 1:200 during the polymerase step, and secondary Abcam donkey anti-mouse Alexa488 antibody (ab150109) was included at 1:500 along with the readout probe in the final hybridization step. mNeon antibody was validated by testing against a non-expressing negative control and a histone-mNeon expressing positive control.

RCA experiments were imaged on a spinning-disk confocal. Z-stacks at 300 nm intervals were captured, and maximum intensity projections were used for all analyses. Hyphae were segmented manually. A Matlab script was used to align hyphae to the tip after manually marking the positions of the tip in each cropped image, marking a second coordinate along the hyphal “axis” and then using the imrotate function to rotate the hypha by an angle calculated as atan(y_axis/x_axis)). Transformed hyphal images were saved and used to create mean projections and to derive radial intensity profiles for Bni1. For Cln3 and cell cycle correlations nuclei were manually assigned cell cycle stages using the Tub1 signal, and RCA amplicons at the nuclear periphery or in the nuclear neighborhood were counted manually, as the spot intensities were too variable across a cell and between hyphae to set an automatic threshold. Cell-cycle-state correlation data was quantified from experiments without *CLN3* smFISH probes, as smFISH spots, while being well colocalized with tri-probes, appeared to compete with them, resulting in lower RCA intensities.

Whi3-IF/RCA images were quantified as follows. Hyphae were segmented manually. Nucleus, RNA and Ribo channels were first subject to a rolling ball background subtraction, and were smoothed with a 1 px Gaussian filter. They were then used as masks after applying a manually corrected Otsu threshold. The nucleus mask was used to obtain the centroid of nuclei. The median Whi3 intensity under RNA masks, and Ribo masks were obtained, and finally the mean ratio of Ribo-associated and RNA-associated intensities were plotted as violin plots in custom Matlab scripts. Cln3 data was additionally binned as Ribo spots lying ≤0.66 µm from nucleus centroids, to determine the translation associated with nuclear-peripheral condensates.
To ensure these colocalizations are not by chance based on the abundance of the punctate signal, non-randomness in spatial distribution of translation relative to Whi3 assemblies was tested by “flipping” the Ribo mask relative to the Whi3 channel. The corresponding “flipped” images showed drastically lower Whi3 intensities indicating these are not just by chance based on underlying Whi3 levels.

| **Probe** | **Sequence** |
| --- | --- |
| Act1Padlock | /5Phos/AAGATA AA CAT CGTAGAC TAGCA TTTCTTTTCTACTGCGTCTAGT CTCAG GTCAT ACACTA |
| Act1Padlock | /5Phos/AAGATA AA CAT CGTAGAC TAGCA TTTGCCACAATGGGTGCAACA CTCAG GTCAT ACACTA |
| Act1Padlock | /5Phos/AAGATA AA CAT CGTAGAC TAGCA ACTCTCTTTAACAGGCACTGTA CTCAG GTCAT ACACTA |
| Act1Padlock | /5Phos/AAGATA AA CAT CGTAGAC TAGCA AGCATAACTAGTTTGGTGGATG CTCAG GTCAT ACACTA |
| BNI1Padlock | /5Phos/AAGATA AA CAT CGTAGAC TAGCA CCGAAAACCTGGGTTGTGTCTCC CTCAG GTCAT ACACTA |
| BNI1Padlock | /5Phos/AAGATA AA CAT CGTAGAC TAGCA TTGGGTCCGAGTTTTACCAGAT CTCAG GTCAT ACACTA |
| BNI1Padlock | /5Phos/AAGATA AA CAT CGTAGAC TAGCA CAGGAGGGTGAGTCTTTAGTC CTCAG GTCAT ACACTA |
| BNI1Padlock | /5Phos/AAGATA AA CAT CGTAGAC TAGCA CCCTGATTTCTGAGTGGGAGG CTCAG GTCAT ACACTA |
| BNI1Padlock | /5Phos/AAGATA AA CAT CGTAGAC TAGCA AACCGGCATCCTCGCTATGTCTT CTCAG GTCAT ACACTA |
| Cln3CDSPadlock | /5Phos/AAGATA AA CAT CGTAGAC TAGCA ATTCCAAAGCGACGATGTTGACC CTCAG GTCAT ACACTA |
| Cln3CDSPadlock | /5Phos/AAGATA AA CAT CGTAGAC TAGCA TTAGGTTCAAGCGTCTGTTTTTCG CTCAG GTCAT ACACTA |
| Cln3CDSPadlock | /5Phos/AAGATA AA CAT CGTAGAC TAGCA AAGGACGAAACTGGGTCGTGA CTCAG GTCAT ACACTA |
| Cln3CDSPadlock | /5Phos/AAGATA AA CAT CGTAGAC TAGCA CATCTGAAAACACGGAGGGTG CTCAG GTCAT ACACTA |
| Act1Primer | GCACGTTGAAAGTTTCAAACCCGTCTACGATG |
| Act1Primer | CGAGAAACCAGCGTAGATAGGCCGTCTACGATG |
| Act1Primer | CGACGTAACATAGCTTCTCTTTAAGTCTACGATG |
| Act1Primer | TCACACTTCATGATGGAGTGAGTCTACGATG |
| BNI1Primer | AGATGGCCCTCAGCTTCAGCCGTCTACGATG |
| BNI1Primer | CCTCACTCTGAAGCTGTCCAGAGTCTACGATG |
| BNI1Primer | GCGCACTACTTTCAAGGTCTCCGTCTACGATG |
| BNI1Primer | CAGAGAATGTGGAGAGGTACAATGTCTACGATG |
| BNI1Primer | CACATTGCGTAGGTTTTCCGAGGGTCTACGATG |
| Cln3CDSPrimer | CCACACATTCCTCCTGATTGAGCC GTCTACGATG |
| Cln3CDSPrimer | TGCAGGCTCGGTATTCGGGTT GTCTACGATG |
| Cln3CDSPrimer | GCGCTGGAGTTGTAATTGTAGTGG GTCTACGATG |
| Cln3CDSPrimer | ATGAGTTGCTCGGAGTAGTTGCC GTCTACGATG |
| Alex647_detection_probe | /5Alex647N/CATACACTAAAGATAACAT |
| Alex555_detection_probe | /5Alex555N/CATACACTAAAGATAACAT |
| Ag18srRNASplint-1 | TAGGGCAGAAATTTGAATGGCATGTAAA AAA AAA AAA AAA AAA AAA AAA AAA AAA AAA AAA AAA AAA AAA AAA AA TATCTTTAGT*G*T*/3InvdT/ |
| Ag18srRNASplint-2 | ACGTCCTATTCCATTATTCCATGCTAAA AAA AAA AAA AAA AAA AAA AAA AAA AAA AAA AAA AAA AAA AAA AAA AA TATCTTTAGT*G*T*/3InvdT/ |
| Ag18srRNASplint-3 | CGATCCCCTAACTTTCGTTCTTGATAAA AAA AAA AAA AAA AAA AAA AAA AAA AAA AAA AAA AAA AAA AAA AAA AA TATCTTTAGT*G*T*/3InvdT/ |
| Ag18srRNASplint-4 | CTGGCCCGTCAGTGTAGCGCGCGTAAA AAA AAA AAA AAA AAA AAA AAA AAA AAA AAA AAA AAA AAA AAA AAA AA TATCTTTAGT*G*T*/3InvdT/ |
| Ag18srRNASplint-5 | GGCCTCACTAAGCCATTCAATCGGTAAA AAA AAA AAA AAA AAA AAA AAA AAA AAA AAA AAA AAA AAA AAA AAA AA TATCTTTAGT*G*T*/3InvdT/ |

### EMSA

In vitro transcribed full length *CLN3* WT or cln3(5m) RNAs were used in 8 reaction conditions including a 0 protein controI. Each 20 µL reactions contained 1X EMSA buffer (20 mM HEPES pH 7.4, 50 µM EDTA, 5% v/v glycerol, 1 mM DTT, 5 mM MgCl2, 10 µg/mL BSA, 150 mM KCl), 1.25 mg/mL *E.coli* tRNA, 0.5 µL murine RNAse inhibitor (NEB M0314S). 2 µL purple DNA loading dye (NEB B7024S), and 5 µL of protein diluted in the Whi3 protein storage buffer () to the desired final concentration. Reactions were allowed to equilibrate at 30ºC for 1 h before running on a 1% agarose gel in TBE. Electrophoresis was carried out at 110 V for 1.5 h. The gels were stained in Sybr Gold for 3 h before imaging.

Gel images were analysed in Fiji. The gel analysis plugin was used to extract intensities of the unbound RNA band across the 8 lanes. The bound fraction was calculated as

Fraction = Int_bound_/[Int_bound_+ Int_unbound_], and ploted against tested protein concentrations.

These curves were fit to the isotherm B_max_*x/(K_D_+x) where x is the protein concentration to estimate upper bounds of the apparent dissociation constant K_d_. Fitting was performed in Matlab 2024 using the fitnlm function.

EMSAs were run in triplicate and K_d_-s for both individual replicates and the fit to the mean binding curve across the replicates were reported.

### Polysome profiling

The polysome profiling protocol for *Ashbya* was adapted from protocols by Valášek (2007) ^8^ and based on Nicchitta lab’s protocols ^9^. Importantly, polysomes were not arrested using cycloheximide, but were crosslinked using 1% formaldehyde.

50 mL of AFM medium was inoculated with 200 µL of dirty spores in 500 mL baffled flasks, followed by incubation at 30°C for 16-17 hours. Whi3-deltapolyQ(6XHAtag)-tdTomato-NAT (AG909) was grown under Nat selection. Cultures were fixed by adding 1.2 mL of cold 37% formaldehyde was added to each 50 mL culture. The cultures were shaken in an ice bath for 5 minutes and then incubated on a 25°C shaker for 10 minutes before being quenched by 5 mL cold 2M glycine.

The fixed cultures were transferred to 15 mL conical tubes, centrifuged at 3000 rpm for 5 minutes, and the supernatant discarded. The cell pellets were resuspended in 700 µL of lysis buffer containing 50 mM HEPES KOH pH 7.4, 140 mM KCl, 1 mM MgCl_2_, 1mM EDTA, 0.1% Tween-20, 5% Glycerol, 4 µL/mL Protease inhibitor cocktail IV (RPI P51000-1), 1mM DTT, 1mM PMSF, 40 U/mL RNAseOUT (ThermoFisher 10777019) and transferred to chilled beadbeater tubes containing 750 µL chilled zirconia beads. Beadbeating was performed for 5 cycles of 90 seconds on and 30 seconds off. The supernatants were transferred to pre-chilled tubes and clarified by centrifugation at 3000g for 5 minutes at 4°C, followed by a second spin at 11,000g for 2 minutes and a third at 11,000g for 10 minutes.

15-50% sucrose gradients were manually prepared by layering a 15% solution over a 50% sucrose solution and the gradient was allowed to form by laying tubes on their side for 2 h. An equal A260 amount of the clarified lysate were carefully loaded into the gradients, not exceeding 1.2 mL. The gradients were centrifuged at 35,000 rpm, 4°C, for 3 hours (L5-50B ultracentrifuge, Beckman). 60% sucrose was used to collect 12x1 mL fractions alongside continuous A254 measurements on a Teledyne Isco fraction collector (Lincoln, NE).

For RNA extraction, polysome fractions were incubated at 56°C for 45 minutes to reverse crosslinking ^10^. RNA was extracted using the Qiagen RNeasy kit (Qiagen 74104). RNA was eluted with RNAse-free water and stored at -80°C.

RNA samples were treated with DNase I, followed by cDNA synthesis using Superscript IV Reverse Transcriptase using a 1:1 oligo-dT and random hexamer primer (ThermoFisher 18090010). The cDNA was stored at -20°C or used directly for qPCR. For qPCR, 3 cDNA dilutions (1x, 0.5x, and 0.25x) were prepared and individual reactions were run for Actin, BNI1 and *CLN3* RNAs using Power SYBR Green master mix (ThermoFisher 4367659), with cycling conditions of 95°C for 2 minutes, followed by 30 cycles of 95°C for 15 seconds and 60°C for 1 minute. Relative cDNA amounts were determined using the ∆Ct method.

| **qPCR primer** | **Sequence** |
| --- | --- |
| act1-1-f1 | CCGTCTTGTCGCTATACTCTTC |
| act1-1-f2 | AACGAGAAACCAGCGTAGATAG |
| bni1-1-f1 | CCAGGAACTTGAACCTGTACTT |
| bni1-1-f2 | GAGTCGCTGAGGTTTCCTATTT |
| cln3-1-f1 | TCCTCCTCGACACTCTTCTTAT |
| cln3-1-f2 | AAGAGCGGTAAGCCCTAGTA |
