## Supplemental Methods for "RNA-specific local translation is patterned by condensates for multinucleate cell growth"

**Model of phase-specific translation**

The bulk translation $T_{bulk}$ is a sum of the contributions of free RNA, RNA associated with Whi3 and dense phase RNA each of which are assumed to show independent translation rates:

$T_{bulk}=k_{1}\Phi_{RNA Free}+ k_{2}\Phi_{RNA-Whi3}+{\epsilon k}_{3}\Phi_{RNA Dense}$	(1)

where ϵ is an RNA-specific factor that determines the translation response in the dense phase. We assume the phenomenological translation rates as Whi3-concentration independent. They can be determined empirically from the luciferase data at three Whi3 concentrations by solving three algebraic equations for the three rates as variables, while setting total RNA concentrations to 5 nM from **Figure 5A,C**. For *CLN3,* at 0 nM Whi3, all RNA is free and total translation is $1*V$, where $V$ is the maximal volume-scaled translation output for a reaction volume $V$. Substituting these values into (1) yields

$V= k_{1}*5 nM*V+0+0, \mathrm{giving} k_{1}=\frac{1}{5}$

At ~150 nM Whi3, which is the measured apparent Kd of the Whi3-*CLN3* complex (**Figure S6**), all RNA is in a single dilute phase with the RNA half-maximally bound. The effective translation is ~$2^{-1}$. Substituting these and $k_{1}$ into (1) and solving for $k_{2}$

$$2^{-1}V= k_{1}*\frac{5}{2} nM*V+k_{2}*\frac{5}{2}nM*V+0, \mathrm{giving} k_{2}=0$$

consistent with steady dilute phase repression as Whi3 concentration increases. At 3 µM Whi3, the system is within the two-phase regime. The measured dilute and dense phase RNA concentrations are 4.95 nM and volume fraction ~97% and 50 nM at ~3%. Additionally, to capture the condensate size-dependent translation (**Figure 7H**), we introduce an inverse radius dependence. Assuming negligible free RNA at this Whi3 concentration, and setting $\epsilon=1$,

$$2^{-10}V= k_{1}*0 nM*V+k_{2}*4.95 nM*V+k_{3}*0.003V*50 nM*\frac{1}{\sqrt{\frac{0.7\mu m^{2}}{pi}}}$$

$\mathrm{gives} k_{3}=5.4*{10}^{-4} \mu m^{2}$.

These rates were introduced into an equilibrium binding and phase separation model where RNA binding to Whi3 in the dilute phase is represented as a saturation curve

$${RNA}_{Free}= \frac{{RNA}_{Total}*{Whi3}_{max}}{K_{d}+{Whi3}_{Total}} \mathrm{nM}$$

where ${Whi3}_{max}= K_{d}=150 nM$.

Dilute and dense phase Whi3 amounts are expressed in terms of the saturation concentration ${Whi3}_{sat}=1000 nM$as

${Whi3}_{Dilute}= {Whi3}_{Total}*\frac{{Whi3}_{sat}}{{Whi3}_{sat}+{Whi3}_{Total}}$, ${Whi3}_{Dense}= {Whi3}_{Total}(1-{Whi3}_{Dilute})$

The dense-phase volume fraction was expressed as a linear function of dense phase protein amount, and was determined empirically using volume fraction at 3 µM Whi3 as

$$V_{Dense}= \frac{{Whi3}_{Dense}*(0.004-0)}{\left( 3000-1000 \right) nM}$$

The dense phase RNA amount was determined using the measured dense phase concentration using the relation

$${RNA}_{Dense}=50 nM*V_{Dense}$$

And mass balance was used to derive dilute phase

$${RNA}_{Dilute}={RNA}_{Total}-{RNA}_{Free}-{RNA}_{Dense}$$

The model produces the expected RNA species distribution as a function of Whi3 (above). Further, the overall translation qualitatively recapitulates the observed luciferase translation (below). Setting $\epsilon=1$ captures the monotonic repression seen in *CLN3,* (red) and increasing the value to 30 shows a peak of translation around the saturation concentration, resembling the translation response of *BNI1*, blue*.* The phase boundary at ${Whi3}_{sat}$ is marked by the dotted red line and reference maximal translation is marked in dotted black line.

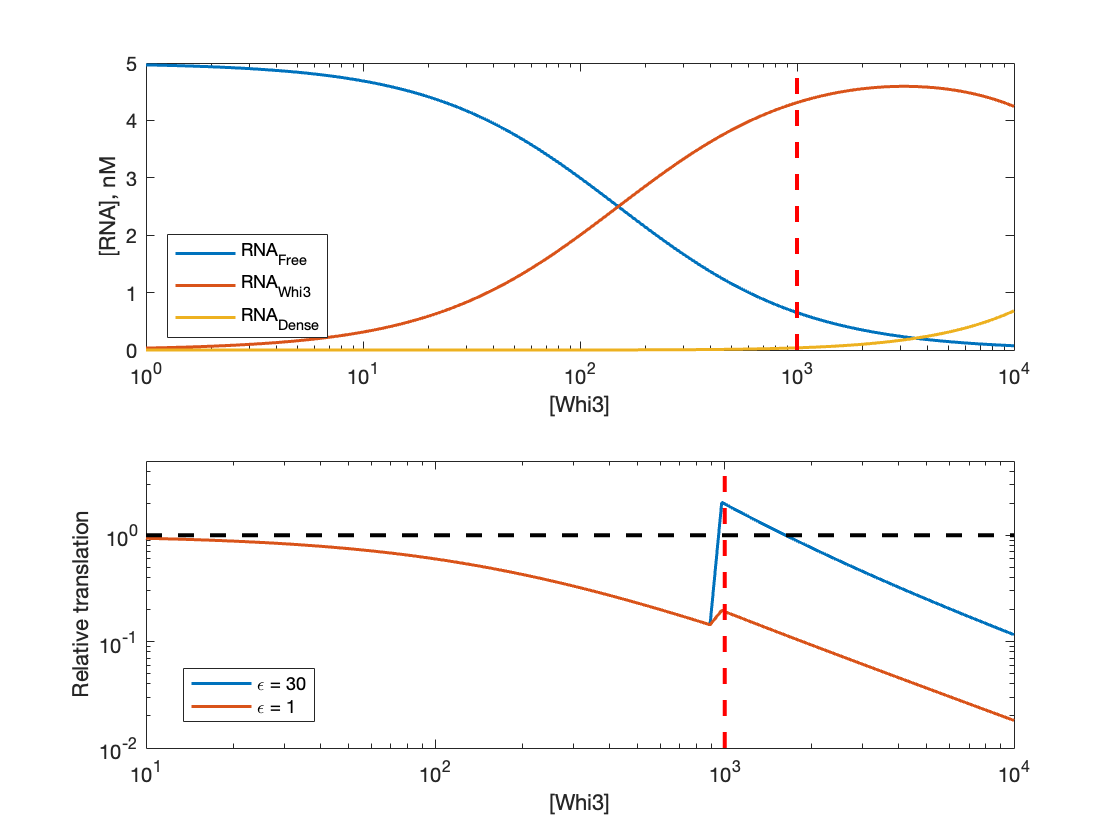
